# Differential impact of GABA_A_ receptors and gephyrin post-translational modifications on layer 2/3 pyramidal neuron responsiveness *in vivo*

**DOI:** 10.1101/2020.11.25.397877

**Authors:** Yuan-Chen Tsai, Mohammad Hleihil, Kanako Otomo, Andrin Abegg, Anna Cavaccini, Patrizia Panzanelli, Teresa Cramer, Kim David Ferrari, Matthew J.P. Barrett, Giovanna Bosshard, Theofanis Karayannis, Bruno Weber, Jillian L. Stobart, Shiva K. Tyagarajan

## Abstract

A diverse set of GABA_A_ receptors (GABA_A_Rs) enable synaptic plasticity adaptations at inhibitory postsynaptic sites in collaboration with the scaffolding protein gephyrin. Early studies helped to identify distinctions between GABA_A_R subtypes allocated within specific functional circuits, but their contribution to the changing dynamics of a microcircuit remains unclear. Here, using the whisker-barrel system in mouse, we assessed the contribution of specific synaptic GABA_A_R subtypes and gephyrin scaffolding changes to sensory processing *in vivo*. We monitored spontaneous and evoked Ca^2+^ transients in layer 2/3 pyramidal cells with the genetically encoded Ca^2+^ sensor RCaMP1.07. Using *Gabra1* or *Gabra2* global and conditional knockout mice, we uncovered that α1- and α2-GABA_A_Rs determine the sparseness of L2/3 pyramidal neuron encoding. In a cell-type dependent manner, α1-GABA_A_Rs and α2-GABA_A_Rs affected neuronal excitability and the reliability of neuronal responses after whisker stimulation. We also discerned that gephyrin with its diverse post-translational modifications (PTMs) shows preference for specific GABA_A_R subtype to facilitate microcircuit activity. Our results underscore the relevance of the diversity of GABA_A_Rs within a cortical microcircuit.

**Key points:**

- While GABAergic inhibition from interneuron subtypes regulates cortical microcircuit activity the molecular determinants have remain unclear.
- We demonstrate that specific-GABA_A_ receptor subtypes contribute differentially to layer 2/3 neuronal activities in mouse barrel cortex.
- Importantly, we link the GABAAR contributions to the scaffolding properties of its important postsynaptic density protein gephyrin. We show that different PTMs on gephyrin determines neuronal excitability via GABAAR recruitment and modulation of inhibition within layer 2/3 neurons.
- Specifically, α1 and α2 subunits containing GABA_A_ receptors, along with their scaffolding protein gephyrin determine the distribution of high, medium and low activity pyramidal neurons during sensory encoding, whereby controlling the total activity of cortical microcircuit.

## Introduction

The primary somatosensory cortex (S1) is composed of excitatory and inhibitory neurons that encode passive sensory experience, training on sensory tasks, and sensory perceptual learning (Feldman and Brecht 2005). Sensory information is processed in a selective manner and is encoded sparsely. The rodent whisker system is a well-established model to study how sensory perception is encoded within organized barrel cortex columns. A whisker deflection leads to depolarization in the majority of neurons, but only around 10% of these neurons respond with action potentials (Crochet et al. 2011). In layer 2/3 (L2/3) of the barrel cortex, the excitability of a cortical microcircuit is tightly modulated by inhibitory interneurons (Kapfer et al. 2007). The excitatory neurons outnumber GABAergic interneurons within the neocortex, although precise numbers vary across different cortical areas and layers. Despite this disparity in excitatory and inhibitory neuron numbers, the probability of finding excitatory-excitatory connections between 2 pyramidal neurons in supragranular layers is relatively low, whereas the connectivity between pyramidal cells and inhibitory interneurons is much higher (Holmgren et al. 2003; Lefort et al. 2009; Avermann et al. 2012; Petersen 2019). Hence, GABAergic interneurons play a crucial role in controlling the activity of the pyramidal neurons within a cortical microcircuit in the superficial layers.

At the postsynaptic sites, GABA_A_ receptors (GABA_A_Rs), its main scaffolding protein, gephyrin, and their associated signaling proteins facilitate downstream signal transduction to modulate pyramidal cell excitability. The importance of GABA for cortical plasticity has been reported for the barrel cortex function, while the molecular mechanisms underlying plasticity changes at GABAergic synapses remain unclear. A local infusion of the GABA_A_R antagonist gabazine leads to increased spontaneous depolarization of L2/3 pyramidal neurons (Gentet et al. 2010). GABA_A_Rs are assembled from a heterogeneous gene pool to form pentameric ion channels. These GABA_A_R subtypes that constitute the GABA_A_R pentamers also exhibit distinct expression patterns within a given brain circuit (Fritschy and Panzanelli 2014). GABA_A_R subtypes are also uniquely localized within different cortical layers. The α1 subunit-containing GABA_A_Rs (α1-GABA_A_Rs) are more uniformly distributed across all six cortical layers, whereas the α2 subunit-containing GABA_A_Rs are abundantly expressed in the supragranular layers (layer 1, 2 and 3), and the α3 subunit containing-GABA_A_Rs are more abundant in layers five and six (Fritschy and Mohler 1995). Hence, this spatial segregation of GABA_A_R subtypes implies that distinct functional roles exist for GABA_A_R subtypes within the somatosensory cortex.

GABA_A_R synaptic localization is facilitated by diverse receptor-interacting proteins that include neuroligin-2 (Fritschy, Panzanelli, and Tyagarajan 2012), collybistin (deGroot et al. 2017; Hines et al. 2018), and gephyrin (Tyagarajan and Fritschy 2014). Among these proteins, gephyrin is unique because it aids activity-dependent adaptations at GABAergic synapses over different time scales (Battaglia et al. 2018). Many signaling pathways converge onto the gephyrin scaffold and in turn induce various post-translational modifications (PTMs) on gephyrin (Zita et al. 2007; Kuhse et al. 2012; Dejanovic and Schwarz 2014; Dejanovic et al. 2014; Ghosh et al. 2016). Super-resolution microscopy studies of gephyrin PTM mutants have shown that compaction of gephyrin molecule and synaptic dwell time of α2 subunit-containing GABA_A_Rs contribute to activity-dependent adaptation (Flores et al. 2015; Battaglia et al. 2018). However, *in vivo* functional significance of gephyrin PTMs on GABA_A_R synapse recruitment and plasticity remains unexplored.

The barrel cortex function is facilicated by GABAergic inhibition via interneuron networks that coordinate pyramidal neuron activity during sensorimotor behavior (Petersen 2014). At the microcircuit level, parvalbumin-expressing (PV+) neurons that typically synapse onto pyramidal cell soma and proximal dendrites become activated during passive and active whisker sensing. In contrast, somatostatin-expressing (SOM+) interneurons that typically synapse onto distal dendrites of a pyramidal cell become hyperpolarized during whisker sensing (Gentet et al. 2012). The importance of inhibition is even more prominent in awake mice as the cortical state changes from quiet to active whisker behavior involves the reorganization of GABAergic neuronal network activity (Gentet et al. 2010). The response of pyramidal neurons to different interneuron inputs during sensory processing relies heavily on the downstream receptor complex and postsynaptic density proteins. Hence, within a cortical microcircuit, differential GABA_A_R activation within the principal cell must contribute to encoding and regulating sensory inputs. Currently, the identities of specific GABA_A_Rs and the role of scaffolding proteins in determining cortical microcircuit functional specificity are unknown.

In this study, we examined the involvement of two major GABA_A_R subtypes, namely α1-GABA_A_R and α2-GABA_A_R, and gephyrin PTMs, for sensory input-dependent cortical microcircuit function. In this regard, we employed a genetically encoded Ca^2+^ sensor, RCaMP1.07, to measure Ca^2+^ transients in L2/3 pyramidal neurons in the barrel cortex in wild-type (WT) and *Gabra1* or *Gabra2* global and conditional gene-deficient mice. We report α1- and α2-containing GABA_A_R subtypes, together with their scaffolding protein gephyrin, facilitate sparse encoding of whisker stimulation-induced sensory response within L2/3 pyramidal neurons. We identify that at a global and a local level, α1-GABA_A_Rs and α2-GABA_A_Rs determine the excitability of L2/3 pyramidal neurons in the opposite manner. Specifically, within L2/3 pyramidal cells, α1 and α2-GABA_A_Rs control sparseness within the responding population. Importantly, distinct gephyrin PTM facilitates inhibitory synapse plasticity via the recruitment of either α1- or α2-containing GABA_A_Rs to synaptic locations. This dynamic process helps fine-tune L2/3 pyramidal neuron excitability. Together, the changing needs of the barrel cortex L2/3 microcircuit during sensory stimulation are facilitated by specific GABA_A_R subtypes and diverse gephyrin PTMs.

## Methods

All experiments were performed in accordance with the European Community Council Directives of November 24, 1986 (86/609/EEC) and approved by the cantonal veterinary office of Zurich.

### Animals

Male and female mice C57BL/6J were purchased (Charles River, Germany), and the following strains were maintained in house: FVB, *Gabra1* KO (Vicini et al., 2001), *Gabra2* KO (Panzanelli et al., 2011; Koester et al., 2013), *Gabra1*^flox/flox^ (Vicini et al., 2001), *Gabra2*^*flox/flox*^ (Duveau et al., 2011). Control groups were sex-matched with the experimental groups. They were group-housed in an inverted 12-hour light/dark cycle. All mice underwent surgery at 8-12 weeks of age and were imaged repeatedly (2-4 times per week) under a 2-photon microscope for up to 5 months.

### Cloning and virus production

The AAV2/6-CaMKIIa-RCaMP1.07 construct was generated by cloning of the RCaMP1.07 gene (Ohkura et al., 2012; Pilz et al., 2015) into an adeno-associated plasmid backbone (AAV2) under a calcium/calmodulin dependent protein kinase II alpha (CaMKIIα) promoter.

The eGFP-gephyrin-K148R, eGFP-gephyrin-DN, eGFP-gephyrin S303/305A expression vectors have been described previously (Flores *et al*., 2015; Ghosh *et al*., 2016), and were subcloned into an AAV2 plasmid backbone containing the human synapsin1 (hSyn1) promoter, in an inverted orientation, and flanked by 2 different loxP sites. The transgene was packaged into AAV 6 serotype. All the AAV6 recombinant viruses were generated by the Viral Vector Core at the University of Zürich. AAV2/6-CaMKIIa-CreER^T2^ virus was purchased from Vector Biolabs (#2014-1208).

### Lateral ventricle viral injection

C57BL/6 mouse pups received bilateral viral injection into lateral ventricle at post-natal day 0 (P0) (Kim et al. 2013). In short, P0 pups were briefly anaesthetized with isoflurane, and 2 μl viral solution per lateral ventricle was injected using a 10 μl Hamilton syringe with a 32 gauge. Lateral ventricles were targeted using bregma and lambda as reference points to draw the mid-line, the needle was inserted into a site 1 mm lateral from the mid-point of the mid-line perpendicular to the skin surface. The needle was inserted to a depth of 3 mm into the skull to ensure injection into the lateral ventricle. Virus used for the experiments was AAV8-CaMKIIa-tdTomato in the control and experimental groups. The injected pups were placed back in their home cage after waking up from anesthesia. The mice were housed under normal conditions until 2-months of age when they were then used for experiments.

### Tamoxifen administration

Tamoxifen (1 mg per animal; Sigma, T5648) was given intraperitoneally (i.p.) for 4 consecutive days to induce Cre recombinase activity from CreER^T2^. The neurons expressing the transgene were imaged 5-7 days post final tamoxifen injection.

### Surgery and virus injections for 2-photon Ca^2+^ imaging

Surgical procedures were divided into 2 steps, which were 2-4 days apart. In the first surgery, after fixing the mouse head in a stable position in a stereotaxic frame, an incision was made along the mid line to expose the skull. After cleaning the bone, bonding reagent (ONE COAT 7 Universal, Coltene) was applied, and then a head cap was created using layers of light-cured dental cement (SYNERGEY D6 Flow, Coltene). Finally, the custom-made aluminium head post was attached to the head cap. These procedures were carried out under isoflurane (4% for induction and 1.5-2% for maintenance, Forene, AbbVie). The second surgery involved a craniotomy, cortical viral injection and chronic window implantation. With a dental drill, a small piece of skull was removed above the sensory cortex to expose the barrel cortex. A glass pipette and hydraulic pump were used to inject 150 nL of virus (injecting speed 50-70 nL per minute) at a depth of 350 μm beneath the brain surface into the whisker areas identified by Intrinsic optical imaging (see below). Immediately after the injections, a 3×3 mm coverslip was fixed right above the exposed brain and secured with dental cement to the head cap. Buprenorphine (Temgesic 0.1 mg/kg) was given before and after surgical procedures for 3 days.

### Intrinsic optical imaging (IOI)

IOI was used to identify barrel areas of the corresponding whiskers in the left somatosensory cortex. This technique was used to image activation of barrel areas through the skull (before craniotomy) to map the whisker field for potential viral injections, and through the cranial window to map specific whisker areas before 2-photon imaging. Under a red light (630 nm illumination), the activated brain region (imaged 400 μm under cortical surface) was identified by increased light absorption following whisker deflection by a piezo stimulator. Imaging was done by using a 12-bit CCD camera (Pixelfly VGA, PCO imaging), and the animals were maintained under 1-1.2% isoflurane.

### Two-photon imaging

A custom-built 2-photon laser-scanning microscope (Mayrhofer et al. 2015) with a 20x water immersion objective (W Plan-Apochromat 20x/1.0 DIC VIS-IR, Zeiss) was used for *in vivo* Ca^2+^ imaging in anaesthetized mice. The microscope was equipped with a Ti:sapphire laser (Mai Tai; Spectra-Physics) set to 1040 nm to excite RCaMP1.07. Fluorescence emission was detected with GaAsP photo-multiplier modules (Hamamatsu Photonics) fitted with a 520/50 nm band pass filter or a 607/70 band pass filter and separated by 560 nm dichroic mirror (BrightLine; Semrock). A customized version of ScanImage (r3.8.1; Janelia Research Campus) was used for setting imaging parameters and to control the microscope.

Confirmation of viral expression at the injected site and neuronal activation during whisker stimulation was examined first, before the series of imaging sessions. Every imaging session comprised 40 trials of spontaneous activity and 40 trials with whisker deflection (90 Hz, 1 s, piezo-based stimulator), while the duration of each trial was 15 s. Fast images were taken (11.84 Hz, 128×128 pixels) to capture neuronal Ca^2+^ responses. Imaging depth ranged from 160-200 μm beneath the cortical surface, which was in layer 2/3 of the barrel cortex. Once a field of view was selected, the same field was imaged for 3-4 sessions on different days, and an extra 3 sessions if the animals were subjected to tamoxifen injections.

### Western Blot analysis

Barrel cortices from both hemispheres were isolated surgically and lysed mechanically in EBC lysis buffer (50 mM Tris-HCl, 120 mM NaCl, 0.5% NP-40, and 5 mM EDTA) containing a protease inhibitor cocktail (cOmplete™, Mini Protease Inhibitor Cocktail, Roche) and phosphatase inhibitors (Phosphatase Inhibitor Cocktail 2, Phosphatase Inhibitor Cocktail 3, Sigma-Aldrich), and then incubated for 1 hour on ice. The lysates were then centrifuged at 23,000 rpm (∼48,500 rcf) for 30 minutes. Samples were prepared for loading by mixing 30 μg of protein in 5xSDS sample buffer containing 15% 2-Mercaptoethanol (Bio-Rad) and heated at 90 °C for 5 minutes. Then samples were loaded onto 8% SDS-polyacrylamide gels and run in Tris-glycine buffer at room temperature. Gels were transferred to a PVDF membrane. The membranes were blocked for 1 hour at room temperature with 5% blocking agent (Roche Diagnostic) in TBST (100 rpm shaker). Primary antibodies in blocking solution were then added to the membrane and incubated overnight at 4 °C (100 rpm shaker). After washing with TBST, the membranes were then incubated for 1 hour at room temperature with secondary antibodies coupled to either horseradish peroxidase or IRDye® (LI-COR) to visualize the protein bands with either film (FujiFilm) or Odyssey imager (LI-COR). Intensity of the bands were quantified using ImageJ. The levels of GABRA1, GABRA2 and GABRA3 were normalized to actin levels.

### Immunohistochemistry

All mice were perfused with ice cold artificial cerebrospinal fluid (ACSF; in mM: 125 NaCl, 2.5 KCl, 1.25 NaH_2_PO_4_, 26 NaHCO_3_, 25 D-glucose, 2.5 CaCl_2_, and 2 MgCl_2_), which was oxygenated (95% O_2_, 5% CO_2_) for 30 minutes. After isolating the brains, the tissues were subjected to 90 minutes post fixation in 4% paraformaldehyde (PFA) at 4°C. The post-fixed brain was left overnight in 30% sucrose at 4°C for cryo-protection. The frozen brains were cut into 40 μm thick coronal sections by using microtome and stored in anti-freeze buffer. Sections within the coverage of the barrel field were selected and stained for the α1 or α2 GABA_A_R subunit (see antibody list) and an appropriate secondary antibody was used to visualize the receptor localization. The images (1024×1024 pixels) were captured using a confocal LSM 700 microscope (Zeiss) and synaptic clusters were analysed using ImageJ image-processing plugin (github repository: https://github.com/dcolam/Cluster-Analysis-Plugin). Z-stack images were taken to cover the cell body and its apical dendrite, and these images were then collapsed into 2D images for cluster analysis. Auto-thresholding moments method was applied. A mask over the somatic region was created for every analysed neuron (10-15 neurons per mouse, 4-5 mice per group) and the mask was enlarged by a factor of 1. Colocalization analysis was then performed for GABRA1 or GABRA2 staining. GABRA1 or GABRA2 particles were identified if the sizes fell within the range between 0.03 μm and 5 μm. Imaging and image processing were performed under blinded condition.

### Slice preparation

For miniature inhibitory postsynaptic current (mIPSC): 4 weeks old mice were injected in the barrel cortex with the described virus. After another 4 weeks, mice were sacrificed by cervical dislocation followed by decapitation under anaesthesia. The brains were removed quickly and immersed in ice-cold cutting solution (in mM): 110 sucrose, 60 NaCl, 3 KCl, 1.25 NaH2PO4, 28 NaHCO3, 0.5 CaCl2, 7 MgCl2, 5 D-glucose. The slices were incubated at 32°C for >30 minutes in ACSF, in mM: 125 NaCl, 2.5 KCl, 1.25 NaH2PO4, 26 NaHCO3, 25 D-glucose, 2.5 CaCl2, and 2 MgCl2. 300 μm coronal slices of the virus-injected area of the somatosensory cortex were made using a Leica VT1200 vibratome (Leica). After the recovery period, slices were maintained at RT with continuous perfusion of carbogenated ACSF (95% O2, 5% CO2). All recordings were performed at RT.

For evoked inhibitory postsynaptic current (eIPSC): virus were stereotactically injected into the L2/3 barrel cortex of 6-8 weeks old mice. After allowing 4 weeks of virus expression, mice were anaesthetized with isoflurane and sacrificed by decapitation and their brains were rapidly harvested and transferred to ice-cold dissecting solution containing (in mM): 110 choline chloride, 35 MgCl2, 25 D-glucose, 25 NaHCO3, 12.5 KCl, 6.25 NaH2PO4, 0.5 CaCl2, saturated with 95% O2 and 5% CO2. 270 μm coronal slices containing barrel cortex were sectioned using a Vibrotome VT 1200S (Leica) then transferred to ACSF containing (in mM): 115 NaCl, 35 KCl, 25 NaHCO3, 25 D-glucose, 12 NaH2PO4, 2 CaCl2, 1.3 MgCl2, continuously aerated with 95% O2 and 5% CO2. Slices were kept at RT and recovered in ACSF for at least 30 minutes before recording.

### Electrophysiology

For mIPSC: Recordings were amplified by Multiclamp 700B amplifier and digitized with Digidata 1440 (Molecular Devices). The recording electrode was pulled from a borosilicate glass pipette (3–5 MΩ) using an electrode puller (PC-100, NARISHIGE Group). Pipettes were filled with caesium-based solution containing (in mM): 120 CsCl, 10 EGTA, 10 HEPES pH 7.4, 4 MgCl2, 0.5 GTP and 2 ATP. Events were isolated by adding CNQX (25 μM, Merck), AP-5 (50 μM, almone labs) and tetrodotoxin (1 μM, Affix Scientific). 10 min after establishing the whole-cell mode, mIPSCs were analysed for a duration of 2 min. Events were recorded using Clampex 10.7 software (Molecular Devices). Recordings were filtered offline using a Bessel low pass filter at 500 Hz (Clampfit 10.7, Molecular Devices) and analysed using MiniAnalysis 6.0.7 software (Synaptosoft). Data were analyzed by ordinary One-way or Brown-Forsythe and Welch One-way ANOVA followed by Tukey’s or Dunnet T3 multiple comparison tests, respectively.

For eIPSC: Slices were placed in the recording chamber of an upright microscope (Axioscope 2 FS, Zeiss) kept at 28.9°C, superfused with ACSF at a rate of 2 ml/min and continuously oxygenated with 95% O2 and 5% CO2. Whole-cell, voltage clamp experiments were performed using borosilicate patch pipettes (1.5 OD X 0.86 ID X 75 L mm, Harvard Apparatus), pulled with a DNZ Universal Electrode Puller (Zeitz-Instruments) to have a resistance of 3-4 MΩ and filled with a solution containing (in mM): 130 CsMeSO3, 10 HEPES, 5 CsCl, 5 NaCl, 2 MgCl2, 2 EGTA, 0.4 GTP, 0.1 EGTA, 0.05 CaCl2 (pH 7.3, 280-290 mOsm/kg). Each cortical pyramidal neuron was voltage-clamped at 0 mV during the recording. IPSCs were evoked by stimulation through a borosilicate glass pipette filled with ACSF, which was connected to a constant current stimulator (Stimulus Isolator Model IS4, Primelec) and placed within 150 μm from the soma of the recorded neuron. A stimulus of increasing amplitude ranging from 4 to 12 uA (0.1 ms, at least 10 repetitions each) was delivered every 10 s to evoke IPSCs.

Data were acquired using an Axopatch 200B amplifier controlled by pClamp software (v10.7, Molecular Devices), filtered at 2 kHz and sampled at 10 kHz (Digidata 1440A, Molecular Device). Series resistance (mean±SEM = 22.8±2.00, range 11–29 MΩ) was monitored at regular intervals throughout the recording and only those recordings with series resistance variations of ≤20% were included in the analysis. Data are reported without corrections for liquid junction potentials. Evoked IPSC data were analysed using Clampfit 10.7 (Molecular Devices). The analysed data had passed normality tests and two-way ANOVA and Sidak’s multiple comparisons tests were performed using Prism 8 (GraphPad).

### Primary neuronal cultures, transfection and immunocytochemistry

Cortical culture on coverslips were prepared from rat (Envigo) embryos at E17. Plasmid transfections were done at 13 days *in vitro* (DIV 13) with 1 μg of total plasmid DNA per coverslip. The plasmid(s) were mixed with 2 μl lipofectamine 2000 (Thermo Fischer Scientific) and 1 μl (1:10 diluted) magnetofection beads (CombiMag, OZ Bioscience). The plasmid-lipofectamine-magnetofection mix were incubated at room temperature for 15 min before adding to neurons. The neuron dishes were placed on top of magnetic plates during the 45 min transfection in the 37°C incubator. The coverslips were transferred to 1 ml conditioned media and returned to the incubator. The plasmids used in the groups are the following: pEGFPC2-gephyrin P1, pEGFPC2-gephyrin K148R, pEGFPC2-gephyrin DN, pEGFPC2-gephyrin S303A/S305A. After one-week of plasmid expression (DIV13+7), the coverslips were rinsed in ice cold PBS, incubated with primary antibody mix containing 10% normal goat serum, GABRA1 (rabbit) and GABRA2 (guinea pig) antibodies was added to the coverslips and incubated for 90 min inside a humidified chamber at 37°C incubator. The coverslips were rinsed in PBS at room temperature (RT), and fixed with 4% PFA for 10 min at RT. After rinsing out the PFA, the coverslips were incubated with corresponding secondary antibodies (see antibody list). Images from ∼20 neurons (3 different batches) were analysed for eGFP-gephyrin clusters colocalization with GABRA1 and GABRA2 containing GABAARs. One-way ANOVA was used to compare between different groups, with post-hoc Tukey test.

### Two-Photon quantification and statistical analysis

All individual neurons expressing RCaMP1.07 in a field of view were manually selected as regions of interest (ROIs) on ImageJ and further processed with custom-designed toolbox Cellular and Hemodynamic Image Processing Suite (CHIPS; Barrett et al., 2018) for MATLAB (R2015b, MathWorks). The images for a field of view were first motion-corrected with a 2D convolution engine for x-y drift. The dF/F value from each manually selected ROI was calculated relative to the baseline imaging period before whisker stimulation (the 2.5 seconds from the beginning of each trial). Peaks were identified by the MATLAB findpeaks function following application of a digital band-pass filter (passband frequency of minimum 0.1 Hz and maximum 2 Hz) and a threshold of 5 standard deviations from the baseline. The peak onset time was calculated by the first time point of the smoothed signal trace (2 frames moving average) crossed over the threshold (the mean of the 2.5 seconds baseline plus 1.5 times the standard deviation) after the start of stimulation.

For the analysis, we focused on the 2 second time window indicated in Fig. 2A (1 second of whisker stimulation and 1 second after the stimulation). For the high responders, the cut-off points for amplitude and number of events were defined as ∼10% of the neuronal population in WT of the corresponding experiment. The size of the population, amplitude, duration and number of events of the high responders were analysed separately.

Statistical analysis was performed using R (version 3.5.3) with the multcomp package for linear-mixed effect model. We set the following as fixed effects which were tested individually or for their interactions: stimulation condition (with or without whisker stimulation), genotype (for *Gabra1* KO and *Gabra2* KO experiments), Cre (for *Gabra1*^*flox/flox*^ and *Gabra2*^*flox/flox*^ experiment, with or without Cre expression), treatment with TAM (for the gephyrin mutant experiment, before or after tamoxifen injections), mutants (for gephyrin the mutant experiment, to compare between different mutants). Individual animal and ROIs were set as random effects. The data were presented with means (un-corrected) and standard error of the means (SEM). The p values reported for different comparisons were obtained by using a Tukey post-hoc test.

### Antibody table

Antibody table:
| Target | Distributor/Produce from | Description & Ref. no. | Dilution | Marker of |
| --- | --- | --- | --- | --- |
| GFP | Aves Labs | Chicken, GFP-1020 | 1:5000 | IHC |
| CaMKIIa | Thermo Fisher Scientific | Mouse, 13-7300 | 1:1000 | IHC |
| GABRA1 | Home-made | Guinea pig | 1:10000 | IHC |
| GABRA1 | Home-made | Rabbit | 1:10000, 1:600 | ICC,<br>WB |
| GABRA2 | Home-made | Guinea pig | 1:2000 | IHC |
| GABRA2 | OriGene | Rabbit, TA327182 | 1:1000 | WB |
| ACTIN | Millipore | Mouse, MAB1501 | 1:10000 | WB |
| GEPHYRIN | Synaptic Systems | Mouse | 1:1000 | WB |

## Results

### GABA_A_R subtypes adapt their expression to altered sensory inputs

To determine whether GABAergic synapses are involved in activity-dependent plasticity in the barrel cortex, we assessed whether GABA_A_R cell surface expression was altered in response to sensory stimulation or sensory deprivation. We injected an AAV8-CaMKIIα-tdTomato into the lateral ventricles of pups on post-natal day 0 (P0) to achieve wide-scale tdTomato expression in excitatory neurons, including cortical pyramidal cells (Kim et al. 2013). At P60, we either stimulated the vibrissae bilaterally for one minute every day or trimmed all the whiskers daily for one week (Fig. 1A). The mice were then sacrificed and tdTomato-positive L2/3 pyramidal neurons in the barrel cortex were analyzed for expression of GABA_A_R subtypes using immunohistochemistry for α1 or α2 subunit containing-GABA_A_Rs, as α1- or α2-GABA_A_R subtypes represent the most abundant synaptic GABA_A_Rs in the superficial cortical layers (Fritschy and Mohler 1995) (Fig. 1B, C). Cluster analysis with an Image-J plugin (see methods) was performed to identify α1- or α2 subunit staining within the somata, where synapses are predominantly inhibitory, and determined the size and density of α1- or α2-GABA_A_R clusters (Fig. 1D-G). The α1-GABA_A_R cluster density was reduced after 1-week of whisker stimulation or whisker trimming (F_(2, 11)_=37.18, P<0.0001; Fig. 1D). In contrast, α1-GABA_A_R cluster size was increased after 1-week whisker stimulation but did not change after whisker trimming (F_(2,10)_=4.6, P=0.036; Fig. 1E). The α2-GABA_A_R cluster density was not changed after either stimulation or trimming (F_(2,10)_=2.27, P=0.153; Fig. 1F), but their cluster size increased in the whisker stimulation group (F_(2,10)_=6.45, P=0.016; Fig. 1G). Together, altered sensory inputs to the barrel cortex led to a dramatic reduction in the number of somatic α1-GABA_A_R clusters, while the α1- and α2 subunit containing-GABA_A_Rs accumulate at inhibitory postsynaptic sites after a week of whisker stimulation. These results indicate that GABA_A_R subtypes adapt differently to changes in sensory input, suggesting differential roles for GABA_A_R receptor subtypes during sensory processing.

**Figure 1.**
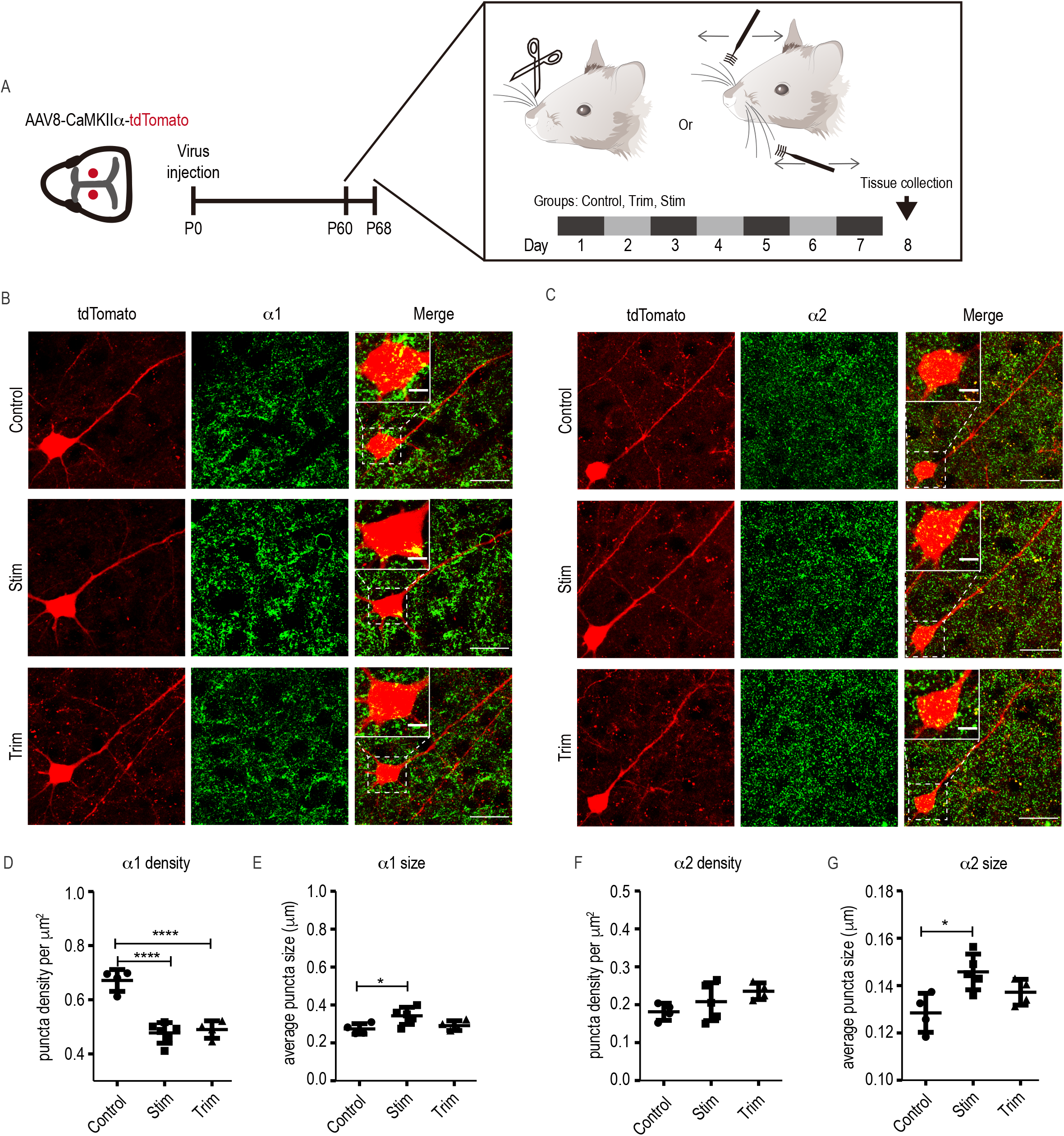
GABA_A_R subunit expression changes after altered sensory input. **(A)** An overview of bilateral viral injections into lateral ventricles at P0, followed by stimulation (Stim) or whisker trimming (Trim) protocol for 1 week prior to morphology analysis on individual pyramidal neurons. Trim group, the whiskers were trimmed bilaterally every day to the same length as the facial hair. The Stim group received daily 1 minute of large angle whisker deflection bilaterally. (B) Representative images of L2/3 pyramidal neurons in barrel cortex from control, trim and stim groups expressing α1-GABA_A_R subunit. (C) Representative images of L2/3 pyramidal neurons in barrel cortex from control, trim and stim groups expressing α2-GABA_A_R subunit. (D-E) Quantification of cluster analysis of density and size for α1-GABA_A_R from N=4 mice for control and trim groups, N=5 for Stim group. (F-G) Quantification of cluster analysis of density and size for α2-GABA_A_R. In total, 10-15 neurons were imaged per mouse and data points represent average of individual animals. Statistics: One-way ANOVA, with Bonferroni post hoc test. Error bar: standard deviation. *p≤0.05,**p≤0.01, ***p≤0.001.

### α1-GABA_A_Rs contribute to sparse pyramidal neuron activity during sensory encoding

The altered expression of GABA_A_R subtypes can potentially lead to changes in neuronal activities, affecting the local microcircuit. To evaluate the functional role of α1-GABA_A_Rs in sensory processing, we made use of the *Gabra1* global gene deletion mouse line (*Gabra1* KO) (Vicini et al. 2001). The activity of the L2/3 pyramidal neurons in the barrel cortex were assessed by using AAV6-CaMKIIα-RCaMP1.07, a red-shifted genetically encoded Ca^2+^ indicator (Ohkura et al. 2012), and *in vivo* 2-photon Ca^2+^ imaging (Fig. 2A). Each neuronal population was imaged three-four times on different days under light anesthesia (1.2% isoflurane). The spontaneous and single whisker stimulation-induced Ca^2+^ transients were imaged and analyzed. In the trials with whisker stimulation, we used a 1-second 90 Hz single-whisker stimulation protocol with a piezo-based stimulator (Mayrhofer et al. 2015; Stobart et al. 2018), and it started 2.5 seconds (baseline) after the trial initiation (Fig. 2A). In order to capture the immediate stimulus-induced Ca^2+^ transients, our analysis was restricted to the time during and after 1 second of whisker stimulation (Figure 2A; right panel). Changes in neuronal activity were assessed by analyzing Ca^2+^ transients for parameters such as amplitude, duration at half-maximum of amplitude, and the number of events per imaging session (40 trials each for spontaneous or whisker stimulation-induced conditions).

**Figure 2.**
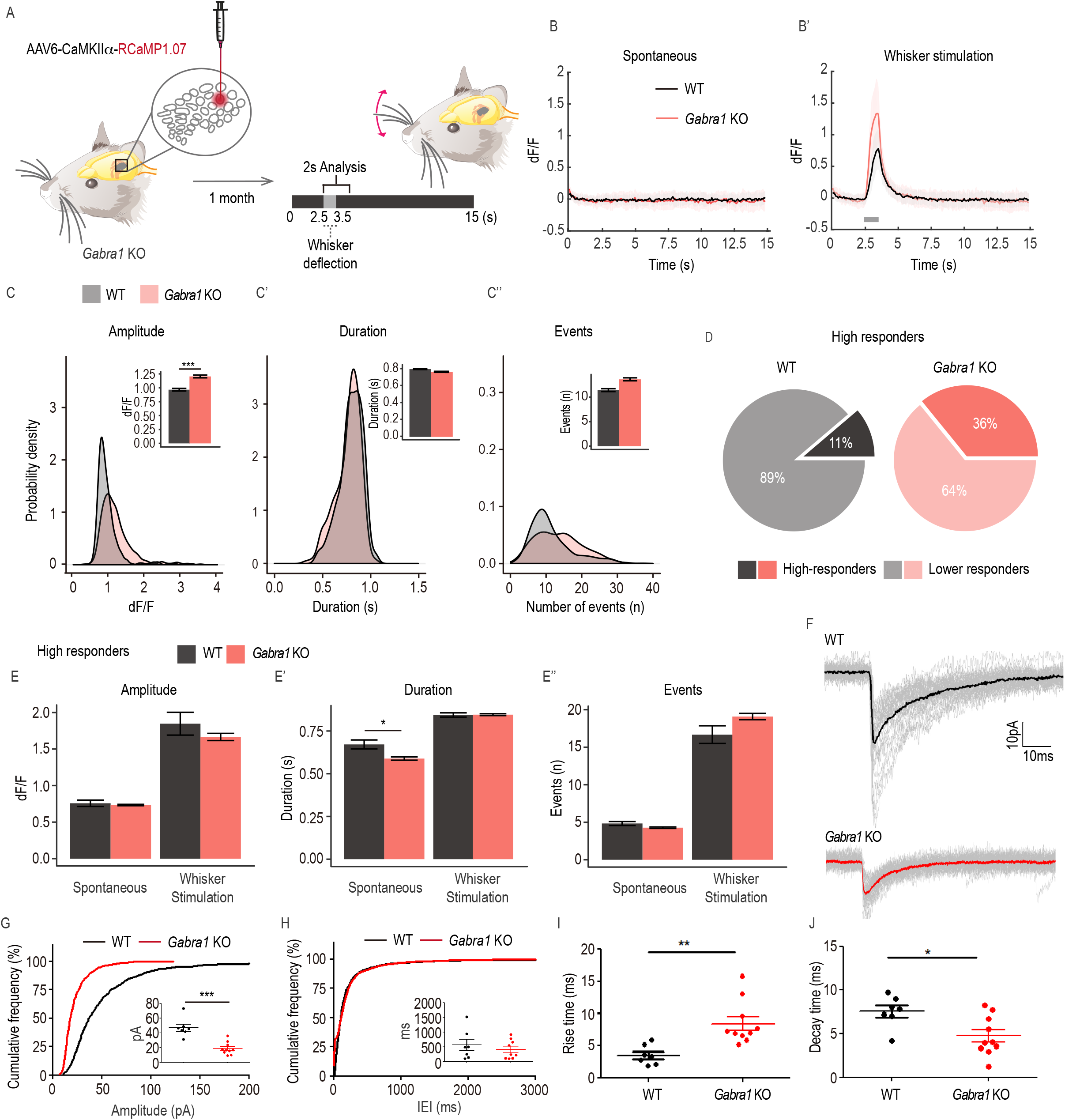
L2/3 pyramidal neuron activity changes in *Gabra1* KO mice. **(A)** Overview of virus injection protocol for *in vivo* 2-photon imaging. AAV6-CaMKIIa-RCaMP virus was injected into the barrel cortex. After 1-month, the mice were subjected to a series of imaging composed of spontaneous and whisker stimulation imaging sessions, and each session was composed of forty 15-second trials (with 3-second inter-trial interval). Each whisker stimulation trial consisted of 1 second of 90 Hz whisker stimulation after 2.5 seconds of baseline. **(B)** Average trace example of spontaneous and whisker stimulation imaging sessions. Grey bar: whisker stimulation. Standard deviations are shown in lighter colors under the example trace. **(C-C’’)** Average Ca^2+^ transient amplitude, duration, and number of events in whisker stimulation trials in density plots and bar graphs (insets). The average was obtained from all imaging sessions for individual neurons (N=459 neurons from 5 *Gabra1* KO mice, N=418 neurons from 4 WT mice). **(D)** Proportions of high and lower responding cells were selected based on ∼10% of high Ca^2+^ transient amplitude versus rest. **(E-E’’)** Quantification of amplitude, duration and the number of events between the high responders of WT and *Gabra1* KO. Statistics: linear mixed-effects models and Tukey post hoc tests. All bar graphs are represented as mean ± SEM. *p≤0.05, **p≤0.01, ***p≤0.001. **(F)** Miniature IPSC (mIPSC) traces were recorded from *Gabra1* WT and KO littermates L2/3 barrel cortex. Average: solid black (WT) and red (KO). All traces are shown in grey. **(G)** mIPSC amplitude **(H)** mIPSC inter-event interval (IEI) from *Gabra1* WT and KO are shown. Cumulative distribution is plotted. Inset: each dot represents a recorded cell (all events averaged). **(I)** Averaged rise time **(J)** and decay time of mIPSC. Each dot represents a recorded cell. Total 3 WT (7 cells) and 3 KO (10 cells). Statistics: Welch’s t-test, mean ± SEM. *p≤0.05, **p≤0.01, ***p≤0.001.

The single-whisker stimulation induced clear Ca^2+^ responses, while the spontaneous Ca^2+^ transients in the same time window demonstrated no change between WT and *Gabra1* KO mice (Fig. 2B; Suppl. Fig. 1B-B’’). However, in *Gabra1* KO animals, L2/3 pyramidal neurons exhibited a higher amplitude of Ca^2+^ transients after whisker stimulation (Fig. 2C). Analysis for the duration and the number of events per neuron (over 40 trials) did not show differences in average values between WT and *Gabra1* KO (Fig. 2C’-C’’). Also, the time of onset and decay time of Ca^2+^ events were similar in WT and *Gabra1* KO (Suppl. Fig. 1C-C’). Pyramidal neuron activity *in vivo* indicates that roughly 10% of the population in the L2/3 barrel cortex encodes sensory stimulation with robust, high amplitude Ca^2+^ events due to sparse firing (Crochet et al. 2011), even though the Ca^2+^ response magnitude is broadly distributed across the population (Fig. 2C). Based on this knowledge, and to understand how *Gabra1* gene depletion impacts the population distribution of pyramidal neuron activity, we defined a cut-off for high-responding cells as the 90^th^ percentile amplitude value from the WT neurons and only considered L2/3 pyramidal neurons in the WT and KO groups that had amplitudes greater than or equal to this value (Margolis et al. 2012). In WT mice, these high responders were 11% of the total population, while high responders with amplitudes over the same cut-off mark were 36% of the *Gabra1* KO principal cells (Fig. 2D). This likely accounted for the higher signal amplitudes observed in the *Gabra1* KO population (Fig. 2C). When we compared the Ca^2+^ transient amplitude, duration and number of events within this subset of high-responding neurons, we did not find any difference between the two genotypes, apart from a reduction in the duration of spontaneous Ca^2+^ transients in *Gabra1* KO mice (Fig. 2E-E’’). Our results show that α1-GABA_A_Rs control pyramidal neuron excitability after whisker stimulation and sparseness within the population.

The thalamus plays a central role in relaying the vast majority of sensory information to the cortex. The α1-GABA_A_Rs expressed within the thalamus have been shown to gate thalamic output to the visual cortex and promote the onset of the critical period of ocular dominance plasticity (Sommeijer et al. 2017). Within the barrel cortex, thalamic afferents innervate mainly L4, with lesser innervations onto L1 and L5 (El-Boustani et al. 2020). To minimize the functional influence originating from the thalamus, we blocked action potentials using tetrodotoxin (TTX) and measured miniature inhibitory postsynaptic currents (mIPSCs) in barrel cortex L2/3 pyramidal neurons of WT and *Gabra1* KO mice. Our electrophysiological recordings showed a significant reduction in the amplitude of GABAergic postsynaptic currents with no changes in the frequency of the events in *Gabra1* KO L2/3 pyramidal neurons (Fig. 2F-H). The *Gabra1* KO L2/3 neurons had faster rise times and slower decay kinetics (Fig.2 I-J). To understand compensatory changes with another GABA_A_R subunit, we used morphology analysis and stained for VGAT, gephyrin, and α2-GABA_A_R subtype in WT and *Gabra1* KO mice cortex (Suppl. Fig. 1D). The cluster analysis showed no changes in VGAT and gephyrin density between WT and *Gabra1* KO, but a significant increase in α2-GABA_A_R density in *Gabra1* KO tissue (Suppl. Fig. 1E-E’’’’). However, the increase in α2-GABA_A_Rs did not fully compensate for the reduced inhibition by the loss α1-GABA_A_Rs. Together, these results emphasize the essential role of α1-GABA_A_Rs for modulating microcircuit activity in response to sensory input in the mouse barrel cortex.

### α2-GABA_A_Rs regulate the reliability of L2/3 pyramidal cells responding to sensory inputs

Next, we evaluated the role of α2-GABA_A_Rs using a *Gabra2* gene global deletion mouse line (*Gabra2* KO) (Panzanelli et al. 2011; Koester et al. 2013). Similar to the approach described above for *Gabra1* KO, spontaneous or stimulation-induced pyramidal neuron Ca^2+^ transients were measured (Fig. 3A-A’). Analysis of spontaneous Ca^2+^ transients showed no genotype effect on amplitude (Suppl. Fig. 2A, B). We observed a wider distribution of signal amplitudes in *Gabra2* KO upon whisker stimulation. Although the mean amplitudes between WT and *Gabra2* KO were not different, the majority of *Gabra2* KO cells had lower than average Ca^2+^ amplitude, and a small sub-group had higher than average amplitude responses (Fig. 3B as an example). The durations of spontaneous Ca^2+^ transients were similar between WT and *Gabra2* KO (Suppl. Fig. 2B’, B’’); however, after whisker stimulation, Ca^2+^ transients tended to have a shorter duration (p=0.13; Fig. 3B’). When considering the number of Ca^2+^ events per neuron across all trials, the frequency of spontaneous activity was similar between *Gabra2* KO and WT cells (Suppl. Fig. 2B’’). Still, whisker stimulation evoked fewer (∼50%) Ca^2+^ transients in *Gabra2* KO, indicating reduced reliability of pyramidal neurons to repetitively encode whisker stimulation (Fig. 3B’’). In addition, we observed slower onset and longer decay time of Ca^2+^ transients after whisker stimulation in *Gabra2* KO compared to WT (Suppl. Fig. 2C-C’).

**Figure 3.**
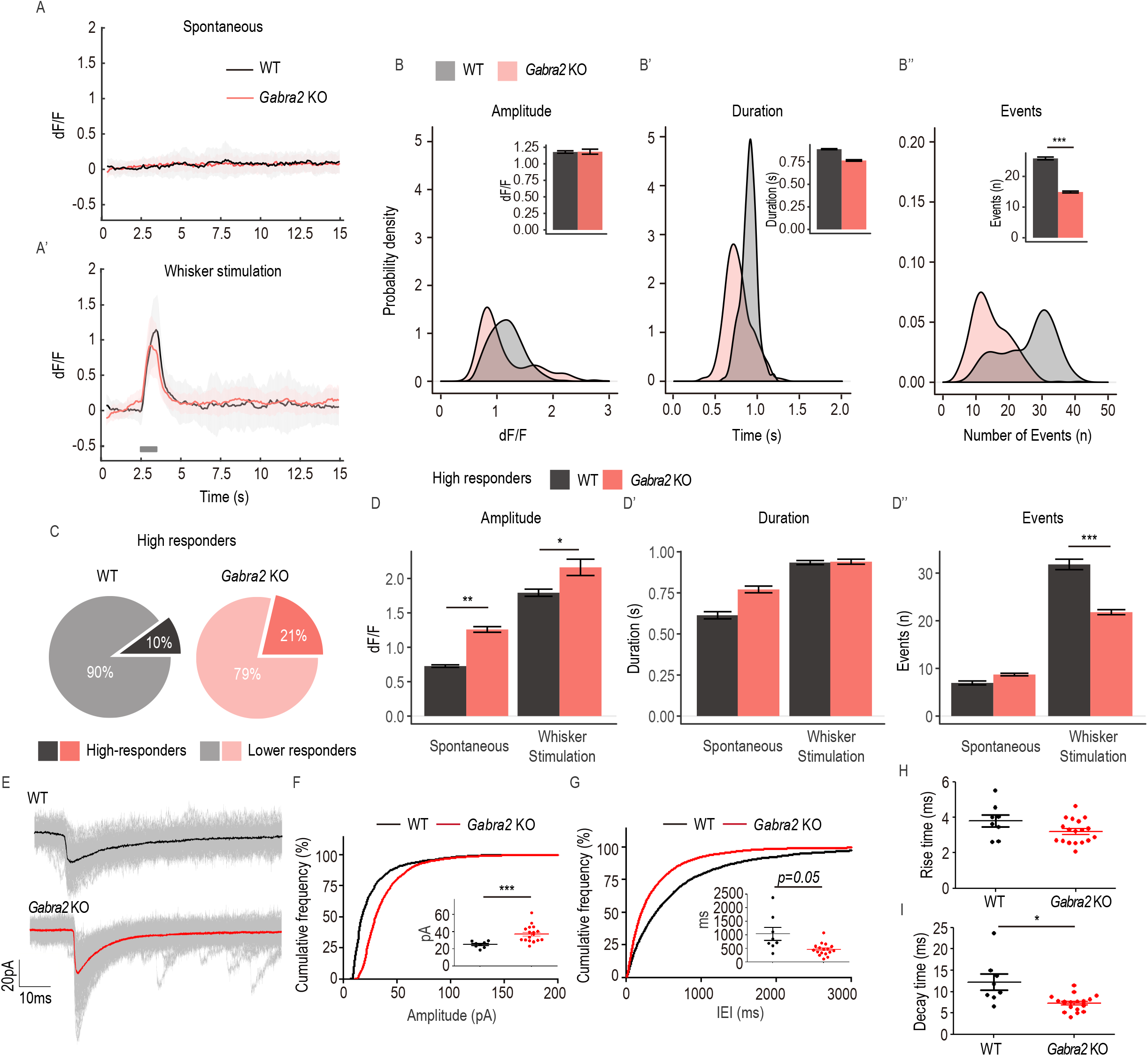
L2/3 pyramidal neuron activity change in *Gabra2* KO mice. **(A)** Average trace from an example of spontaneous and whisker stimulation imaging session. Grey bar: whisker stimulation. Standard deviations are shown in lighter colors below the example trace. **(B-B’)** Average Ca^2+^ transient amplitude, duration and number of events in whisker stimulation trials in density plots and bar graphs (insets). N=370 neurons from 4 *Gabra2* KO mice, N=307 neurons from 3 WT mice. **(C)** Proportions of high and lower responding cells selected based on ∼10% of high Ca^2+^ transient amplitude versus rest. **(D-D’’)** Quantification of amplitude, duration and the number of events measured both high responders in WT and *Gabra2* KO. Statistics: linear mixed-effects models and Tukey post hoc tests. All bar graphs are represented as mean ± SEM. *p≤0.05, **p≤0.01, ***p≤0.001. **(E)** Miniature IPSC traces were recorded from *Gabra2* WT and KO L2/3 barrel cortex. Average: solid black (WT) and red (KO). All traces are shown in grey. **(F)** Amplitude **(G)** and IEI of mIPSC from *Gabra2* WT and KO littermates. Cumulative distribution plotted. Inset: each dot represents a recorded cell (all events averaged). **(H)** Averaged rise time **(I)** and decay time of mIPSC. Each dot represents a recorded cell. Total 3 WT (8 cells) and 4 KO (17 cells). Statistics: Welch’s t-test, mean ± SEM. *p≤0.05, **p≤0.01, ***p≤0.001.

To evaluate whether *Gabra2* gene deletion also influences the population distribution of highly responsive pyramidal neurons, we set a cut-off according to the amplitudes of the WT group, as described in the previous section. Using this amplitude value, we found that 21% of pyramidal neurons were high responders in *Gabra2* KO mice (Fig. 3C). Within this subset of high responding cells, the amplitude of both spontaneous and whisker stimulation-induced Ca^2+^ transients in *Gabra2* KO was higher than in WT high-responders (Fig. 3D). At the same time, there was no difference in the duration of spontaneous or whisker stimulation-induced events (Fig. 3D’). Although the number of events per neuron occurring under spontaneous activity was not different between high amplitude pyramidal neurons of each genotype, the number of events evoked by whisker stimulation was reduced in *Gabra2* KO (Fig. 3D’’). Our data suggest that α2-GABA_A_Rs influence the success rate of whisker stimulation-triggered neuronal responses and partially contribute to the sparseness of the high-responding pyramidal neurons by increasing the neuronal activity in a small set of the neuronal population.

To evaluate how α2-GABA_A_Rs contribute to the Ca^2+^ transients of L2/3 pyramidal neurons, we blocked action potentials using tetrodotoxin (TTX) and measured miniature inhibitory postsynaptic currents (mIPSCs) in barrel cortex L2/3 pyramidal neurons of WT and *Gabra2* KO mice. Our electrophysiological recordings showed increased mIPSC amplitude in *Gabra2* KO (Fig. 3E, F), and the inter-event intervals were reduced (Fig. 3G). The rise time for receptors did not change in the *Gabra2* KO, while the decay kinetics was reduced (Fig. 3H, I). The observed changes in electrophysiological properties within *Gabra1* KO and *Gabra2* KO suggest that compensatory changes at synaptic sites are different in the absence of α1 or α2 subunit-containing GABA_A_Rs. We stained for VGAT, gephyrin, and α1-GABA_A_R subtype in WT and *Gabra2* KO mice cortex (Suppl. Fig. 2D). The cluster analysis showed no changes in VGAT, gephyrin, or α2-GABA_A_R density between WT and *Gabra2* KO (Suppl. Fig. 2E-E’’’’).

To understand compensatory changes of GABA_A_R subunits at the protein level, we used Western blot analysis to assess barrel cortex tissue and measured expression changes in α2 and α3 subunits in *Gabra1* KO and α1 and α3 subunits in *Gabra2* KO (Suppl. Fig. 3). As shown previously (Kralic et al. 2006), expression of the α3 subunit was increased in *Gabra1* KO with no change in α2 subunit levels (Suppl. Fig. 3A, A’). In *Gabra2* KO tissue, we also found an increase in the α3 subunit expression but no change in α1 subunit expression (Suppl. Fig. 3B, B’). The increased expression of α3, however, is unlikely to account for observed effects on Ca^2+^ transients in the *Gabra1* and *Gabra2* KOs.

### Cell-autonomous effects of GABA_A_R subtypes within barrel cortex microcircuits

While our results strongly suggest that GABA_A_R subtypes are important for modulating microcircuit activity and contribute to function differentially, the compensation from other GABA_A_R subtypes in the global KO models appears to be prominent. Hence, we moved to the *Gabra1*^*flox/flox*^ or *Gabra2*^*flox/flox*^ mice to conditionally ablate the *Gabra1* or *Gabra2* gene from adult L2/3 pyramidal neurons using cell-type-specific Cre expression. We injected AAV6-CaMKIIα-RCaMP1.07 or co-injected AAV6-CaMKIIα-eGFP-Cre and AAV6-CaMKIIα-RCaMP1.07 into the barrel cortex and compared Ca^2+^ transients between RCaMP only cells (control) and Cre-positive RCaMP cells (Fig. 4A). Post-mortem validation of the deletions of GABRA1 and GABRA2 in *Gabra1*^*flox/flox*^ and *Gabra2*^*flox/flox*^ mice was done by immunohistochemistry (Suppl. Fig. 4A-B)

**Figure 4.**
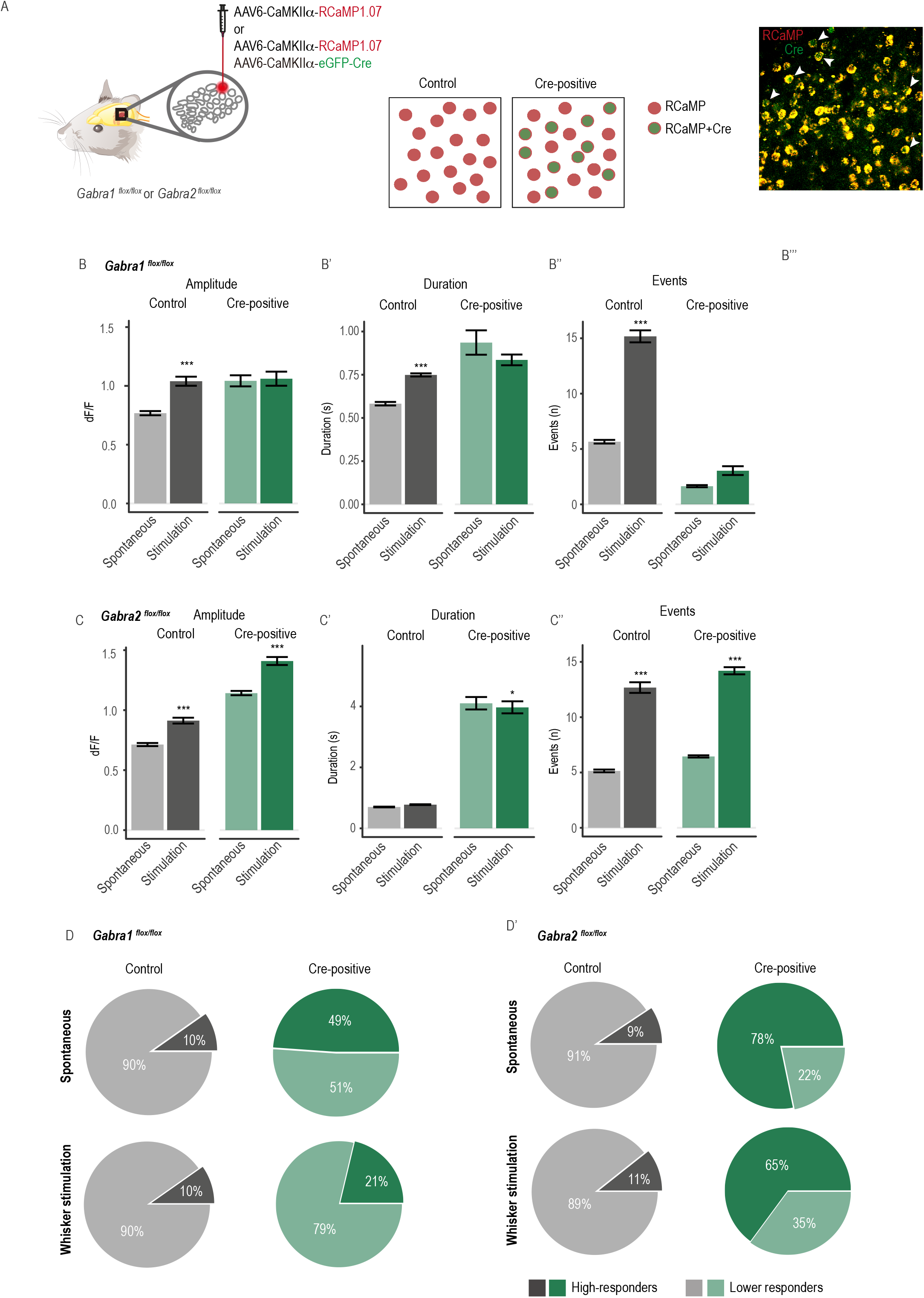
Cell-autonomous contribution of α1-GABA_A_R and α2-GABA_A_R in L2/3 pyramidal neuron whisker derived activity. **(A)** Method illustration. The *Gabra1*^*flox/flox*^ or *Gabra2*^*flox/flox*^ mice were injected with AAV6-CaMKIIα-RCaMP (control) or combined with AAV6-CaMKIIα-eGFP-Cre. The cell population received the viral mixture RCaMP and Cre, named Cre-positive (red cells with green nuclei). Right panel: representative picture from a field of view expressing RCaMP and Cre. White arrowhead: example of cre-positive cells. **(B-B’’)** Spontaneous and stimulation-induced Ca^2+^ transient amplitude, duration, and the number of events for control and Cre-positive RCaMP-expressing neurons from *Gabra1*^*flox/flox*^ mice. N=123 control neurons and N=47 cre-positive neurons from 2-4 mice per group. **(C-C’’)** Spontaneous and stimulation-induced Ca^2+^ transient amplitude, duration, and the number of events for control and Cre-positive RCaMP-expressing neurons from *Gabra2*^*flox/flox*^ mice. N=150 control neurons and N=262 Cre-positive RCaMP-neurons from 3-5 mice per group. **(D-D’)** Proportions of high- and lower-responding cells selected based on 90^th^ percentiles of the control group for spontaneous and stimulation-induced Ca^2+^ transient amplitude in *Gabra1*^*flox/flox*^ **(D)** and *Gabra2*^*flox/flox*^ mice **(D’)**. The same amplitude cut-off levels are then applied to the control and Cre-positive neurons of *Gabra1*^*flox/flox*^ and *Gabra2*^*flox/flox*^ mice. Statistics: linear mixed-effects models and Tukey post-hoc tests. All bar graphs are represented as mean ± SEM. *p≤0.05, **p≤0.01, ***p≤0.001, the stars indicate the significance of the comparison between spontaneous and stimulation. The significance of cross-group comparisons are indicated in the text.

Next, we quantified changes in Ca^2+^ transients in *Gabra1*^*flox/flox*^ mice (Fig. 4B-B’’; Suppl. Fig. 4C-E). The RCaMP control pyramidal neurons showed an apparent increase in Ca^2+^ amplitude to whisker stimulation. Surprisingly, the Cre-positive RCaMP cells did not show an increase in Ca^2+^ amplitude upon whisker stimulation (Fig. 4B). Similarly, the Cre-positive neurons showed no change in duration and number of events. Interestingly, the number of events was largely reduced in the cells with Cre-expression (Fig.4B’’; Spontaneous: control vs. cre-positive, p=0.1616; Whisker stimulation: control vs. cre-positive, ***p<0.001). To understand whether the local microcircuit is compromised in the absence of α1-GABA_A_Rs, we analysed the Cre-negative RCaMP neurons next to the Cre-positive RCaMP neurons. Our analysis of Ca^2+^ transients in the Cre-negative cells showed similar results as Cre-expressing neurons upon whisker stimulation (Suppl. Fig. 4E). This indicates a subpopulation of neurons lacking α1-GABA_A_Rs can influence the local microcircuit activity more strongly than one had anticipated. Together, our data identify that in the absence of α1-GABA_A_Rs, L2/3 pyramidal neurons exhibit reduced excitability, which contrasts with our global *Gabra1* KO data.

We used the same approach to assess Ca^2+^ transient changes in *Gabra2*^*flox/flox*^ mice (Fig. 4C-C’’; Suppl. Fig. 4F). The effect of removing α2-GABA_A_R could be assessed by comparing RCaMP control with Cre-positive RCaMP cells. Upon whisker stimulation, Ca^2+^ amplitude and the number of events significantly increased in all groups (Fig. 4C, C’’). However, the baseline amplitude in Cre-positive RCaMP areas was higher (control vs. Cre-positive, *p=0.012). Similar to *Gabra1*^*flox/flox*^ animals, we analyzed Cre-negative RCaMP cells neighboring the Cre-positive cells and uncovered that they showed a tendency towards the extended duration of Ca^2+^ transients (Suppl. Fig. 4F middle panel). In contrast to the global *Gabra2* KO results, conditional deletion of *Gabra2* in L2/3 pyramidal neurons reduces inhibition. Furthermore, showing an increase in Ca^2+^ transient amplitude and number of events in Cre-negative RCaMP cells upon whisker stimulation suggests that removing α2-GABA_A_Rs also functionally contributes to reciprocal connections within L2/3 pyramidal cells and their role in maintaining local circuit dynamics.

We further examined the subpopulation of high-responding neurons in spontaneous and whisker stimulation-induced conditions in *Gabra1*^*flox/flox*^ or *Gabra2*^*flox/flox*^ mice (Fig. 4D-D’). In *Gabra1*^*flox/flox*^ mice, 50% of Cre-positive group exhibited high-responding population in spontaneous activity. However, only 21% of high-responding cells had increased Ca^2+^ transients after whisker stimulation. In *Gabra2*^*flox/flox*^ mice, 80% of Cre-positive neurons exhibited higher spontaneous activity compared to 10% in the control neurons. This representation of high-responsive cells remained at 65% after whisker stimulation. In summary, our results demonstrate that α1- and α2-GABA_A_Rs in L2/3 pyramidal neurons have distinctive roles in controlling neuronal excitability at the individual level and local microcircuit level. While removal of α1-GABA_A_R subtypes dampens excitability of the L2/3 circuit, removal of α2-GABA_A_R subtypes facilitates excitability of the circuit.

### Gephyrin scaffold dynamics influence GABAergic neurotransmission in vivo

GABA_A_Rs synapse recruitment via lateral mobility on the plasma membrane, receptor insertion at extrasynaptic sites, internalization, and degradation of synaptic receptors are all dynamically facilitated by gephyrin and various PTMs on it (Tyagarajan and Fritschy 2014). Hence, it is logical that we also ascertain the *in vivo* relevance of this scaffolding protein for α1- or α2-GABA_A_R function within the barrel cortex. Of the various PTMs reported for gephyrin, phosphorylation at Ser 303 and Ser 305 sites is known to be activity-dependent (Flores et al., 2015). It has been reported that phosphorylation of S303 and S305 residues by protein kinase A (PKA) and calcium calmodulin kinase II α (CaMKIIα) respectively facilitates NMDA receptor-dependent GABA_A_R recruitment at inhibitory terminals (Flores et al. 2015). Similarly, SUMOylation at Lys 148 is known to stabilize gephyrin scaffold and GABAergic synapses (Ghosh et al. 2016). Therefore, we selected phosphorylation-null gephyrin mutants S303A/S305A (SSA) to block the activity-dependent recruitment of GABA_A_Rs to postsynaptic sites. Alternatively, we selected the K148R gephyrin mutant to stabilize GABA_A_Rs at synaptic sites. A SUMO1 conjugation-defective mutant (K148R) has been reported to stabilize sub-membrane gephyrin clusters at inhibitory postsynaptic sites (Ghosh et al. 2016). In addition, we used the dominant negative mutant (DN) lacking the last 12 amino acids to disrupt gephyrin scaffolding and GABAergic neurotransmission *in vitro* (Ghosh et al. 2016) (Fig. 5A).

**Figure 5.**
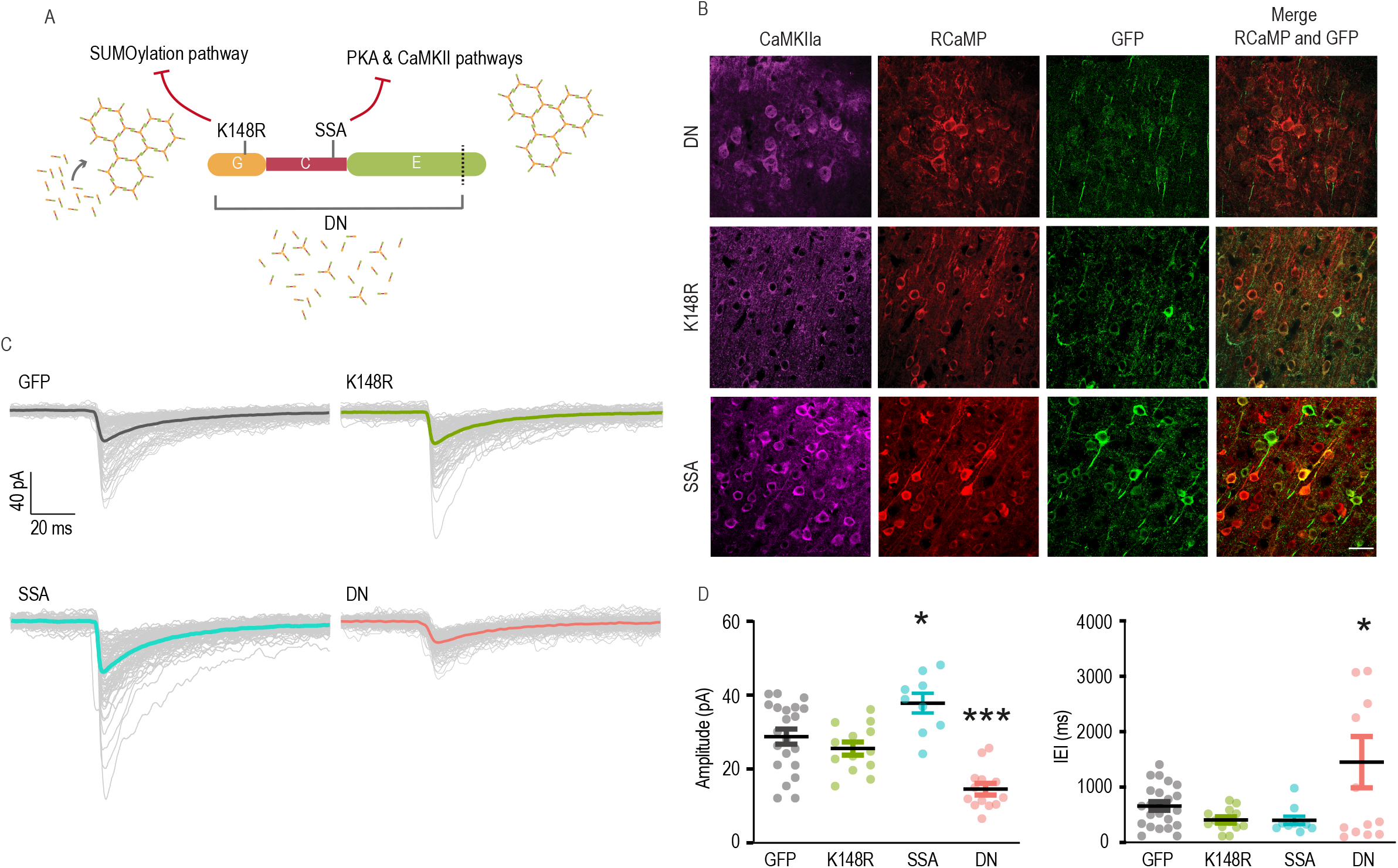
GABAergic neurotransmission changes in gephyrin-mutant-expressing L2/3 pyramidal neurons. **(A)** An illustration of different gephyrin mutations used in the study, along with the signaling pathways that are affected by the mutated residue. The gephyrin-K148R (SUMO1 conjugation site mutant) facilitates scaffolding. The gephyrin-S303A and S305A (PKA and CaMKIIα phospho-null mutant) hampers NMDA receptor activity-induced scaling at GABAergic postsynaptic sites. The gephyrin-DN (lacks part of E domain) disrupts endogenous gephyrin scaffolds in neurons. **(B)** Representative images of neuron co-expressing RCaMP and eGFP-gephyrin variants after injection of AAV in L2/3 barrel cortex *in vivo*. The following combination of viruses were injected to the barrel cortex (same combination used in 2P Ca^2+^ imaging): AAV6-hSyn1-flex-gephyrin mutant, AAV6-CaMKIIa-CreER^T2^, AAV6-CaMKIIa-RCaMP1.07. All brain sections were stained for CaMKIIα and eGFP. **(C)** Example traces from L2/3 pyramidal cells expressing gephyrin variants or GFP control. Colored traces: averaged current from one neuron. Grey traces: all events measured from one neuron. **(D)** Miniature inhibitory postsynaptic currents (mIPSCs) amplitude and the inter-event interval (IEI) in L2/3 pyramidal neurons expressing individual gephyrin variants. Statistics: eGFP, n=23; K148R, n=13; gephyrin-DN, n=13; gephyrin-SSA, n=10 neurons, from 4-5 mice in each group. One-way ANOVA with Tukey post-hoc test. Bars: mean ± SEM. *p≤0.05, **p≤0.01, ***p≤0.001, ****p≤0.0001.

The functional impact of a static gephyrin scaffold or the absence of a gephyrin scaffold at GABAergic postsynaptic sites in L2/3 pyramidal neurons was characterized. For this, we injected AAV6-hSyn1-flex-eGFP-gephyrin variants, AAV6-CaMKIIα-ERT2-Cre and AAV6-CaMKIIα-RCaMP1.07 to co-express eGFP-gephyrin variants and RCaMP1.07 in L2/3 principal cells. This combination of viruses was also used to evaluate activity changes in pyramidal neurons with *in vivo* 2-photon Ca^2+^ imaging (see the next section). After four weeks of virus co-expression, we injected tamoxifen intraperitoneally (i.p) for four consecutive days to activate Cre recombinase and allowed the expression of gephyrin mutant variants (Fig. 5B). We then waited for a least seven days and then recorded miniature inhibitory postsynaptic currents (mIPSC) from L2/3 pyramidal neurons overexpressing the respective gephyrin transgenes in acute slices of barrel cortex, in the presence of tetrodotoxin (TTX), CNQX and AP-5 (Fig. 5C). Neurons expressing the transgene gephyrin-DN demonstrated reduced mIPSC amplitude (14.6 ± 1.6 pA) with respect to GFP controls (28.8 ± 2.0 pA; P<0.0001). The mIPSC inter-event interval (IEI) increased in gephyrin-DN neurons, indicating the frequency was reduced (inter-event intervals: 1451 ± 463.2 ms; GFP=659.9 ± 78.3 ms; P=0.038; Fig. 5D). The distribution of IEI for DN is wider, so we further examine its lognormality by performing D’Agostino-Pearson test, in addition to QQ-plot (Suppl. Fig. 5). The results suggest that the distribution of IEI for DN follows the normal distribution. It is known that gephyrin-DN destabilizes synaptic GABA_A_Rs *in vitro*, and perhaps a similar mechanism reduces the availability of synaptic GABA_A_R in gephyrin-DN-expressing neurons. The neurons expressing gephyrin-SSA mutation demonstrated mIPSCs of increased amplitude (37.8 ± 2.7 pA; P=0.026; Fig. 5D), likely as a result of increased GABA_A_R retention at synaptic sites (Battaglia et al. 2018). Finally, the expression of gephyrin transgene with the K148R mutation had no effect on mIPSC amplitude compared to controls (28.8 ± 2.0 pA vs. 25.6 ± 1.8 pA; P=0.641). The inter-event interval (IEI) was not different between the gephyrin mutants (eGFP: 659.9 ± 78.3 ms; gephyrin-K148R: 408.0 ± 56.3 ms; gephyrin-SSA: 401 ± 73.7 ms), suggesting that there were no major changes in spontaneous neurotransmitter release or in the number of GABAergic inhibitory synapses (Fig. 5D).

Overall, our results suggest that the majority of the GABAergic postsynaptic sites containing gephyrin are influenced by cellular signaling events that directly impact PTMs on gephyrin. Importantly, different gephyrin mutants influence GABAergic neurotransmission in a mutually exclusive manner.

### Gephyrin mutant expression influences whisker stimulation-induced Ca^2+^ transients

As gephyrin scaffold can impact GABAergic neurotransmission in an opposite manner, we measured changes in Ca^2+^ transients in pyramidal neurons before and after gephyrin-mutant expression. Gephyrin-mutant expression was controlled by tamoxifen-inducible Cre expression. We confirmed that around 40% of the pyramidal cells co-expressed all three viral vectors: AAV6-hSyn1-flex-eGFP-gephyrin variants, AAV6-CaMKIIα-ERT2-Cre and AAV6-CaMKIIα-RCaMP1.07 (Suppl. Fig. 6). As an additional control, we expressed RCaMP1.07 in a neighboring barrel area (Area 1) (Fig. 6A). We examined all RCaMP-expressing neurons and compared changes in Ca^2+^ transients before and after eGFP-gephyrin expression. Examples of average Ca^2+^ transient changes in response to whisker stimulation both before and after gephyrin-mutant expression are shown (Fig. 6B). The activity of individual neurons was normalized to activity before tamoxifen (TAM) injection [(After-Before)/Before] to identify relative changes.

**Figure 6.**
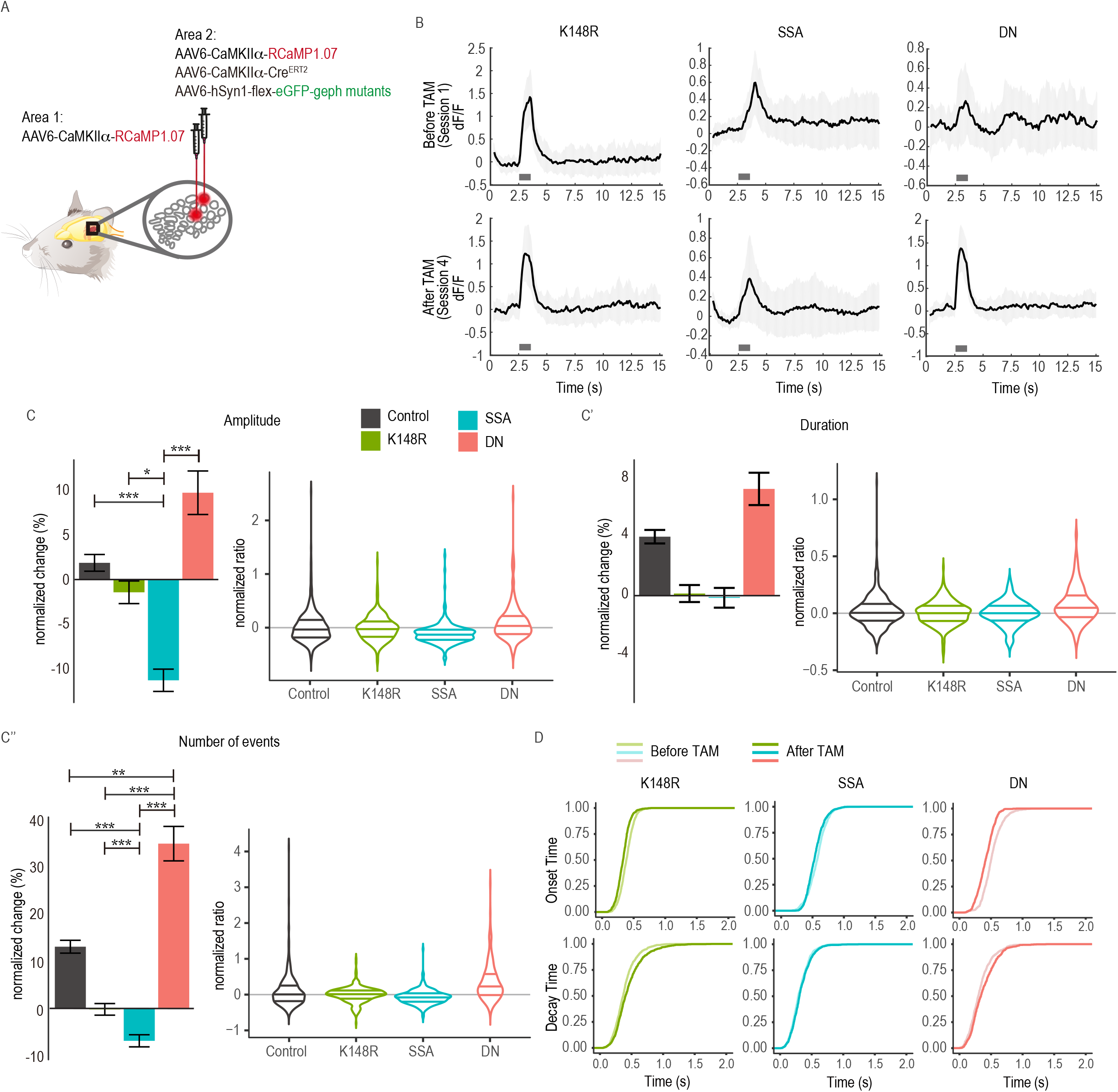
The expression of gephyrin mutants modulate L2/3 pyramidal neuron excitability differentially. **(A)** An overview of the viral infection *in vivo*. The control site (Area 1) received only AAV6-CaMKIIα-RCaMP1.07, while the experimental site (Area 2) received a combination of RCaMP/CaMKIIα-eGFP-Cre-gephyrin viruses. **(B)** Average trace examples from imaging sessions for whisker stimulation trials of both before and after gephyrin mutant expression. Grey bar: whisker stimulation. Lighter colors around the example trace are standard deviations. **(C-C’’)** Average percentage changes after gephyrin-mutant expression for individual neurons are shown in bar graphs and the ratios are shown as violin plots to demonstrate the distribution differences in stimulation-induced Ca^2+^ transients. **(D)** Onset and decay time for before and after expression of the gephyrin mutants. eGFP, n=491 neurons; gephyrin-K148R, n=308 neurons; gephyrin-SSA, n=249 neurons; gephyrin-DN, n=204 neurons. Statistics: linear mixed-effects models and Tukey post hoc tests. All bar graphs are represented as mean ± SEM. *p≤0.05, **p≤0.01, ***p≤0.001.

The gephyrin-mutant expression did not influence spontaneous Ca^2+^ transient amplitude and number of events compared to the control (area1) (Suppl. Fig. 7A, A’’). However, gephyrin-K148R mutant expression reduced spontaneous Ca^2+^ duration (Suppl. Fig. 7A’). Upon whisker stimulation, the gephyrin-K148R mutant caused no change in any of the parameters (Fig. 6C-C”). The gephyrin-SSA mutant expression decreased the amplitude of evoked events, while the gephyrin-DN mutant increased the amplitude of Ca^2+^ transients (P=0.373; Fig. 6C). The duration of Ca^2+^ transients was mostly unaffected by the expression of either of the gephyrin mutants (Fig. 6C’). Expression of the gephyrin-SSA and gephyrin-DN mutants reduced and increased the number of events, respectively (Fig. 6C’’). Furthermore, onset and decay times were similar before and after the expression of gephyrin variants (Fig. 6D). The influence of gephyrin mutant variant expression on Ca^2+^ transient changes was consistent with their impact on inhibitory neurotransmission changes observed using the patch clamp technique (Fig. 5).

**Figure 7.**
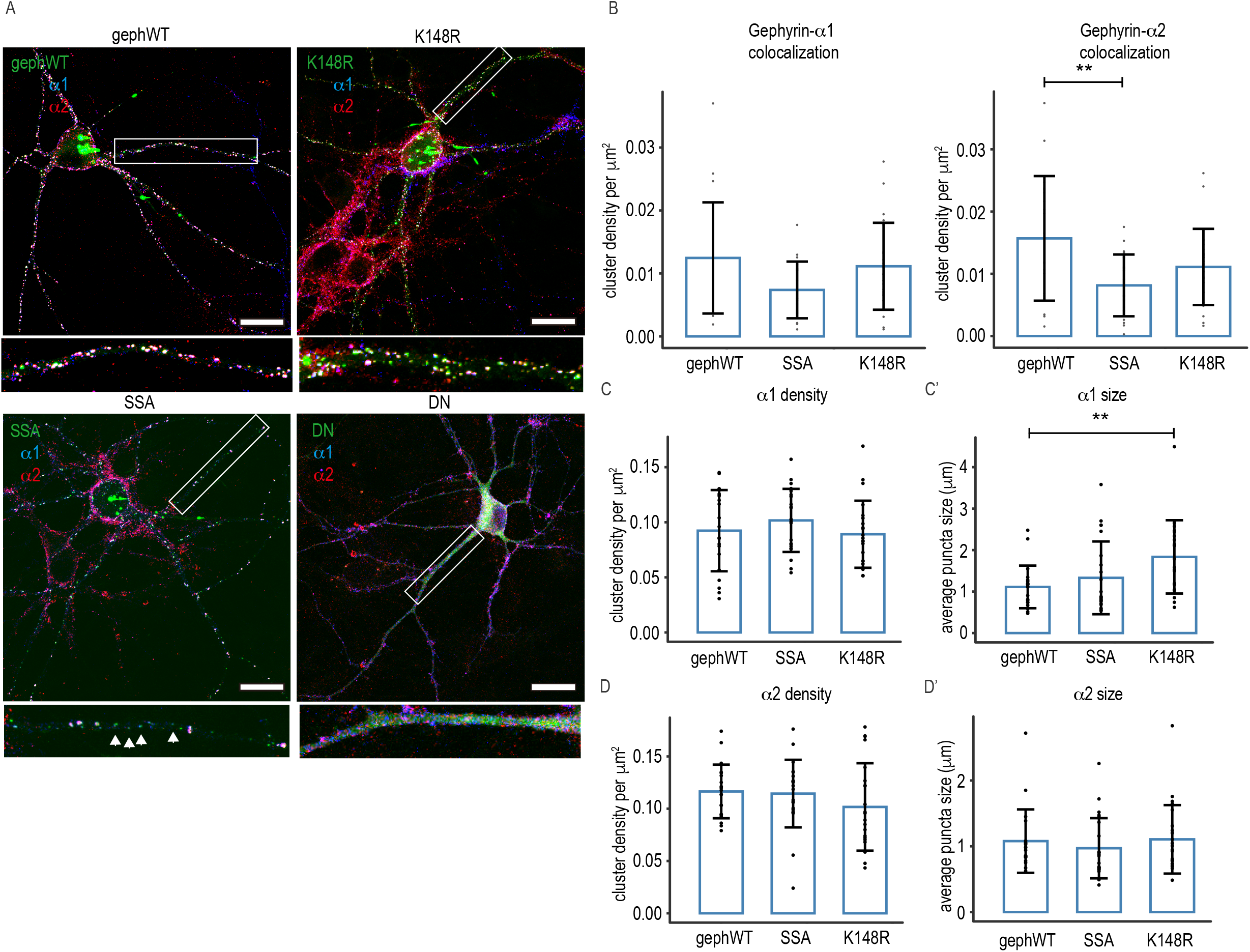
Interaction of gephyrin variants and α1- or α2-GABA_A_R subtypes. The primary cortical neurons were transfected with eGFP-gephyrin WT, eGFP-gephyrin DN, eGFP-Gephyrin K148R or eGFP-gephyrin SSA mutant at 13 days in vitro (13 DIV) and stained for α1 and α2 subunit of GABA_A_Rs at 20 DIV. (A) Example images of transfected neurons with gephyrin mutants. Scale bar: 25 μm. Lower panels: magnified images of selected dendrites (white box). White arrows: eGFP-gephyrin clusters without GABRA2 staining. (B) Cluster analysis was performed to show the density of colocalization of eGFP-gephyrin variants with α1-GABA_A_Rs and α2-GABA_A_Rs. eGFP-gephyrin DN was excluded from cluster analysis as the GFP signal was diffused. Statistics: One-way ANOVA, with Tukey post hoc test. *p≤0.05, **p≤0.01, ***p≤0.001.

We previously identified high responding cells in both conditional and global KO of α1- and α2-GABA_A_R subunits (Figs. 2-4). We used a similar strategy to group neurons based on their Ca^2+^ transients’ amplitude or the number of events within our gephyrin mutant populations (Suppl. Fig. 7B-C). As expected, in control neurons (Area 1), the amplitude and number of events within the populations remained comparable before and after tamoxifen injection. Similarly, gephyrin-K148R mutant expression did not impact the population distribution based on amplitude changes (Suppl. Fig. 7B). However, expression of gephyrin-SSA and gephyrin-DN mutants had the most impact on amplitude changes within the given population of neurons. Specifically, gephyrin-SSA mutant expression led to a smaller population of neurons responding with high amplitude. In contrast, gephyrin-DN mutant expression increased the population of high amplitude neurons by 20% (Suppl. Fig. 7B). The number of Ca^2+^ transient event distributions was not affected upon gephyrin-K148R or gephyrin-SSA expression but was vastly increased in gephyrin-DN mutant expressing L2/3 pyramidal neurons (Suppl. Fig. 7C). Specifically, the cell population exhibiting 21-30 events increased from 29% to 49% and 31-40 events increased from 4% to 10%. The effects of gephyrin mutants on mIPSC amplitudes and frequencies are consistent with the Ca^2+^ transients measured in pyramidal neurons expressing individual gephyrin mutants.

To confirm gephyrin mutation has direct functional impact on L2/3 pyramidal neuron excitability, we recorded stimulation-evoked IPSCs from gephyrin-DN expressing neurons. For this, we infected AAV8-mCaMKIIα-mCherry-2A_iCre and AAV6-hSyn-flex-eGFP or AAV6-hSyn-flex-eGFP-gephyrin-DN were stereotactically injected into L2/3 of the barrel cortex. In both conditions, the co-expression of mCherry and eGFP allowed the identification of co-infected pyramidal neurons. We found that the input/output curve of the eIPSC amplitude at different stimulus intensities for gephyrin-DN mutant was significantly reduced as compared to eGFP control specifically when the stimulus intensity was high. (Suppl. Fig. 5C, D). A two-way ANOVA of eIPSCs revealed that the interaction between the effects of the gephyrin virus and stimulus intensity was statistically significant (F (8, 56) = 3.745, P=0.0014). The results suggest that the manipulation of gephyrin scaffolds by gephyrin-DN overexpression leads to an activity-dependent reduction of inhibition on L2/3 pyramidal neurons in the barrel cortex. Overall, our data identify the gephyrin scaffold as an essential component for response reliability to sensory inputs, and modulations of the gephyrin scaffold in a subset of pyramidal neurons were sufficient to fine-tune the population response to whisker stimulation.

### Gephyrin mutants show a preference for GABA_A_R subtypes

*In vitro* studies have identified gephyrin scaffold as a signaling hub at GABAergic postsynaptic sites to facilitate activity-dependent synaptic GABA_A_R recruitment and removal. However, it remains unclear how specific PTM on gephyrin alters its ability to recruit either α1- or α2-containing GABA_A_Rs. To understand this better, we used cortical primary neuron culture and examined the co-localization of α1 or α2 subunits with eGFP-gephyrin mutants. We transfected primary cortical neurons with either eGFP-gephyrin (WT), eGFP-gephyrin K148R, eGFP-gephyrin SSA, or eGFP-gephyrin DN at 13 days in vitro (13 DIV). At 20 DIV, the cultures were co-stained for α1 and α2 GABA_A_R subunits (Fig. 7). The eGFP-gephyrin or its mutant variants were co-labelled with α1 and α2 GABA_A_Rs, except the DN mutant that showed diffuse signal across soma and dendrites and α1, α2 GABA_A_Rs were also diffusely labeled in DN expressing neurons. The eGFP-gephyrin and its mutants exhibited phenotypes consistent with earlier reports (Flores et al. 2015; Ghosh et al. 2016). Our cluster analysis to determine the colocalization of eGFP-gephyrin or its mutants with either α1 or α2 GABA_A_Rs showed differential preference. The colocalization of WT, SSA, and K148R mutant gephyrin with α1 containing GABA_A_Rs was not significantly different (Fig. 7B). However, SSA mutant exhibited significantly reduced colocalization with α2 containing GABA_A_Rs (Fig. 7B’). The overall density of α1 GABA_A_Rs was not altered in neurons expressing either WT or mutant eGFP-gephyrin (Fig. 7C), but the neuron expressing K148R mutant gephyrin had a significantly bigger α1 GABA_A_R cluster size (Fig. 7C’). The cluster density and size of α2 GABA_A_Rs were similar in neurons expressing either eGFP-gephyrin WT, SSA, or K148R (Fig. 7D, D’). Our results highlight the significance of different post-translational modifications on gephyrin for the recruitment of specific GABA_A_R subtypes to facilitate cortical circuit function.

## Discussion

Somatosensory information is processed in a selective manner and is encoded sparsely. While it is well established that GABAergic inhibition plays a central role in sparse sensory processing, the contribution of GABA_A_R subtypes to the functional specificity of the somatosensory cortex microcircuit is unknown. To fill this knowledge gap, we present data from three groups of cells from floxed *Gabra1* mice or floxed *Gabra2* mice: RCaMP in naïve animals (controls), RCaMP analysis from Cre+ L2/3 pyramidal neurons, and RCaMP analysis from Cre-L2/3 pyramidal neurons (cells neighboring the Cre+ ones). This data set provides the unique opportunity to study the conditional deletion of alpha subunits (Cre +ve cells) vs. control and changes that occur in other nearby pyramidal cells (Cre -ve cells) in the same microcircuit.

Our results from these three groups underscore the contribution of α1- and α2-containing receptors for intrinsic pyramidal neuron excitability and the microcircuit function (which involves pyramidal-pyramidal-GABAergic interneuron interconnectivity to form a local circuit). We also report that at the morphological level, neuronal expression of α1- and α2-GABA_A_Rs are differentially altered after sensory stimulation and deprivation.

We link the functional contributions of α1- or α2-GABA_A_Rs to their main scaffolding protein gephyrin, and its dynamic scaffolding configurations facilitated by PTMs. Specifically, we show that sites S303/S305 phosphorylated in consequence to glutamatergic synapse activity reduces gephyrin colocalization with α2-GABA_A_Rs. Similarly, expression of gephyrin-SSA mutant *in vivo* had the most impact on amplitude changes within L2/3 neurons after whisker stimulation. Specifically, gephyrin-SSA mutant expression led to a smaller population of neurons responding with a high amplitude of Ca^2+^ transients (Suppl. Fig. 7B). On the other hand, stabilizing gephyrin scaffold using K148R mutation facilitated the recruitment of more α1-GABA_A_Rs to synapses. *In vivo* expression of gephyrin-K148R mutant had no impact on the population distribution based on amplitude changes (Suppl. Fig. 7B). By linking the gephyrin mutant study with global and cell-specific conditional *Gabra1* or *Gabra2* KO, our results provide the first direct evidence linking gephyrin scaffold dynamics to GABA_A_R selectivity and L2/3 pyramidal neuron responses upon sensory inputs.

### Neuronal network activity defined by GABA_A_R subtypes

We provide evidence that global ablation of α1-GABA_A_Rs increases pyramidal cell excitability and dampens L2/3 pyramidal neuron inhibition. On the other hand, L2/3 pyramidal neuron-specific deletion of *Gabra1* reduces excitability at both single cell and microcircuit levels. Why would the conditional loss of an inhibitory receptor subunit lead to a decrease in neuronal responses (at rest and during sensory stimulation)? In other aspects of our study, we found similar results where neuronal responses and the number of events were reduced in two other cases. First, in *Gabra2* KO mice, there was a decrease in the number of Ca^2+^ events in the whole population and an increase in the mIPSC amplitude measured by electrophysiology (Fig. 3). Furthermore, there was an increase in GABRA3 expression (Supp. Fig. 3), indicative of an upregulation of other inhibitory subunits, which may account for the observed changes in Ca^2+^ and mIPSCs. Second, expression of gephyrin mutant SSA also decreased the number of Ca^2+^ events (Fig. 6C”) and increased the mIPSC amplitude (Fig. 5), which may occur as a result of increased GABA_A_R retention at synaptic sites (Battaglia et al. 2018). In both of these cases, the results suggest an increased inhibitory tone on pyramidal cells, possibly due to the upregulation of other subunits. In the case of *Gabra1* floxed mice, the precise mechanism of this change remains to be investigated in future studies by electrophysiology and histology for *Gabra* subunit clusters, but it suggests that there is an imbalance in the inhibitory activity that occurs in the whole circuit (both cells with the *Gabra1* deletion and their neighbors). Other *Gabra* subunits (such as *Gabra3*) could be elevated as compensation.

The global ablation of α2-GABA_A_Rs reduces pyramidal cell excitability, microcircuit activity and increases inhibition in L2/3 pyramidal neurons. The L2/3 pyramidal neuron-specific deletion of *Gabra2* increases pyramidal cell excitability and microcircuit activity. In addition, α2-GABA_A_Rs contribute to the reliability of neuronal responses to whisker stimulation. When *Gabra2* subunits are conditionally deleted (Cre + cells), there is a drastic increase in Ca^2+^ activity. Both Ca^2+^ amplitude and the duration of events increase during spontaneous activity and whisker stimulation (Fig. 4C, C’). This suggests that inhibition of pyramidal cells is lost, leading to increased firing that translates to larger Ca^2+^ events.

Interestingly, this elevation in neuronal activity also impacts nearby neighboring pyramidal cells that do not have *Gabra2* deletion (Cre-cells). This suggests that the loss of *Gabra2* was probably not functionally compensated by other alpha subunits. Further, this emphasizes the reciprocal connections within L2/3 circuits and their role in maintaining circuit dynamics. Future studies will help to disentangle the mechanisms behind these changes in *Gabra2* floxed mice.

Barrel cortex plasticity is a multifaceted process involving multiple synaptic and cellular mechanisms. Our data posits that GABA_A_R subtypes are well placed to regulate various aspects of synaptic plasticity mechanisms to recruit silent neurons into the active population. Whether any specificity exists between a GABA_A_R subtype and a subgroup of interneurons is highly debatable. It has been proposed that specific GABA_A_R subtypes are allocated to distinct interneuron terminals, thereby defining a functional circuit. For example, it has been suggested that α1-GABA_A_Rs are localized at PV+ basket cell terminals, while α2-GABA_A_Rs are localized at CCK+ cell terminals (Nyíri, Freund, and Somogyi 2001; Freund 2003). However, it was subsequently reported that α1-and α2-GABA_A_R subtypes could be localized at both CCK+ and PV+ terminals (Kerti-Szigeti and Nusser 2016). While the α-subunit has been long thought to be an essential factor determining the allocation of GABA_A_Rs to different postsynaptic sites, it was recently reported that GABA_A_Rs containing β3 subunits are allocated specifically to PV+ terminals but not SOM+ connections (Nguyen and Nicoll 2018). Hence, there are multiple molecular determinants underlying GABA_A_R synapse allocation, and these factors add an additional layer of complexity to understanding circuit activity.

In hippocampal pyramidal neurons, α2-GABA_A_Rs are preferentially expressed at the axon-initial segment (AIS) when compared with α1- or α3-containing GABA_A_Rs (Nusser et al. 1996; Muir and Kittler 2014). In the somatosensory cortex, this compartmentalized distribution of GABA_A_R subunits is less clear, as α1, α2, and α3 subunits exhibit a similar distribution ratio between the AIS compartment and non-AIS compartment (Gao and Heldt 2016). Subcellular localization differences between GABA_A_Rs are also influenced by lateral surface mobility in response to network activity changes, thereby contributing to synaptic scaling mechanisms (Bannai et al. 2009). Specifically, at AIS, α2-GABA_A_Rs show lower mobility compared to α1-GABA_A_Rs (Muir and Kittler 2014). Therefore, the abundance of α2-GABA_A_Rs at synapses might facilitate membrane depolarization to a greater extent than other GABA_A_R subtypes. It has been reported that stimulating GABAergic axo-axonic cells can elicit action potentials in the postsynaptic pyramidal neurons from L2/3 somatosensory cortex (Szabadics et al. 2006). Hence, α1- and α2-GABA_A_Rs at AIS are well positioned to facilitate pyramidal neuron excitability.

### GABA_A_R subtypes facilitate sparse encoding in L2/3 pyramidal cells

Sparse coding within pyramidal neurons is determined by changes in excitation-inhibition balance (Andermann and Moore 2006). The L2/3 pyramidal neurons have more hyperpolarized resting potentials; therefore, they require more excitatory input and/or reduced GABAergic input to overcome the action potential threshold (Lefort et al. 2009). Our data demonstrate that GABA_A_R subtypes within L2/3 pyramidal neurons are well placed to contribute to circuit-specific sub-threshold excitation and sparse action potential firing. Using *Gabra1*^*flox/flox*^ and *Gabra2*^*flox/flox*^ mice, we demonstrate that a greater proportion of pyramidal neurons become high-responders upon whisker stimulation. In our experiments, we present data from three groups of cells from *Gabra1*^*flox/flox*^ and *Gabra2*^*flox/flox*^ mice: RCaMP in naïve animals (controls), RCaMP analysis from Cre-positive L2/3 pyramidal neurons, and RCaMP analysis from Cre-negative pyramidal neurons (neighboring Cre-positive cells; Suppl. Fig.4 E, F). This data set allowed us to study the effects of conditional deletion of α1 or α2 subunits of GABA_A_Rs (Cre+ cells) vs control, but also changes that occur in other nearby pyramidal cells (Cre-cells) in the same microcircuit. Our results from these three groups highlight the contribution of α1 and α2 subunit-containing GABA_A_Rs towards intrinsic pyramidal neuron excitability and also microcircuit function (which involves pyramidal-pyramidal-GABAergic interneuron interconnectivity to form a local circuit).

In our experiments, we obtained up to 40% co-expression of eGFP-gephyrin mutants with RCaMP and Cre within the L2/3 pyramidal neurons. In spite of this small-scale co-infectivity ratio, we successfully perturbed microcircuit function upon gephyrin-mutant expression in the barrel cortex. Importantly, we showed a role for cellular signaling and gephyrin scaffold PTMs in modulating the size of responding neuronal population to single-whisker stimulation. Disruption of the gephyrin scaffold through the expression of the gephyrin-DN-mutant led to an expansion in neuronal populations with higher amplitude in Ca^2+^ transients. On the other hand, expression of the gephyrin-SSA mutant led to a decrease in the number of high responder cells (Suppl. Fig. 7). Although a population of gephyrin-dependent and independent GABA_A_Rs co-exist, the gephyrin-containing inhibitory synapses functionally dominate and facilitate dynamic shifts in Ca^2+^ transient amplitude adjustments (Suppl. Fig. 7B).

### GABA_A_Rs and gephyrin-mediated homeostatic adaptations within L2/3 microcircuit

It is currently hypothesized that both Hebbian and non-Hebbian synaptic plasticity mechanisms might underlie the strengthening and weakening of cellular responses after whisker stimulation and/or trimming (Turrigiano and Nelson 2000; Hofer et al. 2011; Crochet et al. 2011; Margolis et al. 2012). We observed both strengthening and weakening of pyramidal neuron activity, depending on the α1- or α2-GABA_A_R deletion or gephyrin mutant expression, suggesting that both receptor and its scaffolding protein contribute to homeostatic plasticity mechanisms within cortical L2/3 microcircuit. For example, the ratio of high responders that encoded the whisker stimulus increased in *Gabra1* and *Gabra2* global or conditional KO mice, while the number of Ca^2+^ transients during stimulation decreased (Fig. 2, 3, 4). Furthermore, we found an increase in the mIPSC amplitude in global *Gabra2* KO mice (Fig. 3F). This suggests that the excitation of the pyramidal neuronal network is scaled down in cases of *Gabra2* global KO and *Gabra1* pyramidal neuron-specific KO, even though an inhibitory receptor subtype is lost. These observed changes in pyramidal neuron activity are also reflected upon gephyrin mutant expression *in vivo*, suggesting that homeostatic plasticity mechanisms at GABAergic synapses cannot be completely uncoupled from the receptor and its scaffolding protein.

The gephyrin scaffold is known to interact with α1-GABA_A_Rs and α2-GABA_A_Rs (Tyagarajan and Fritschy 2014). Independent quantum dot–based single-GABA_A_R tracking studies have shown that the gephyrin scaffold influences GABA_A_R surface mobility and synapse retention (Choquet and Triller 2013). Specifically, it was recently shown that gephyrin PTMs, specifically the gephyrin-SSA mutant, increase the synaptic confinement of α2-GABA_A_Rs in response to 4-Aminopyridine-induced circuit activation (Battaglia et al. 2018). Supporting this finding, our electrophysiology data demonstrated an increase in mIPSC amplitude in gephyrin-SSA expressing neurons (Fig. 5), and *in vivo* imaging data showed that a less excitable population of neurons doubled in size upon gephyrin-SSA expression and whisker stimulation (Suppl. Fig. 7B). Increased synaptic confinement of GABA_A_Rs in gephyrin-SSA expressing neurons could disrupt dynamic shifts in reducing inhibitory synapses according to excitatory inputs in a neuron. On the other hand, it has been demonstrated that gephyrin-DN expression increased diffusion and exploration of α2-GABA_A_Rs at both synaptic and extrasynaptic sites (Battaglia et al. 2018). In our electrophysiology studies, we observed increases in mIPSC IEI and reduced amplitude upon gephyrin-DN mutant expression (Fig. 5). *In vivo* imaging data demonstrated that gephyrin-DN mutant expression increases the population of high responders after whisker stimulation by 20% (Suppl. Fig. 7B). Our data implicate a dynamic recruitment model for specific GABA_A_R subtypes based on gephyrin PTM. This would mean that specific demand would trigger a signaling cascade to converge onto the gephyrin scaffold. The PTM change on gephyrin in turn would switch affinity towards specific GABA_A_R subtype for synapse recruitment.

Independent of gephyrin scaffold modulation, a mechanism involving microRNA miR376c was recently reported to specifically regulate dendritic α1 and α2 subunit expression after inhibitory long-term potentiation (iLTP), but not α3, α4, α5 subunit containing-GABA_A_Rs (Rajgor et al. 2020). This highlights a role for multiple context-dependent GABA_A_R modulations at inhibitory synapses.

In summary, while it is known that the distribution of activity within cortical neurons is sparse, our data demonstrate that pyramidal neuron excitability is defined by GABA_A_ receptor subtypes and facilitated by gephyrin scaffold dynamics. Our results have characterized the first-time molecular components that are operational at GABAergic postsynaptic terminals to refine and define the sparse sensory encoding process in adult mice.

## Acknowledgements

We thank J-M. Fritschy for comments on the manuscript and M. Müller for comments on the electrophysiology data. The study was supported by UZH Forschungskredit Can-doc grant to Y-C. Tsai, Olga Mayenfisch Foundation grant, Swiss National Science Foundation grants (31003A_159867 and 310030_192522) and University of Zurich internal funding to S.K. Tyagarajan. Fig. 4 was partially created using BioRender.

## Author contributions

SKT and YCT conceptualised the study. SKT, YCT and JLS contributed to the design of experiments. YCT performed all 2-photon imaging experiments, JLS aided the data analysis. YCT wrote the first draft of the manuscript. SKT, BW, JLS, KDF and MJPB provided technical knowledge and data analysis support. MH carried out electrophysiological experiments and the analysis. KO, AA conducted surgeries for virus injections. KO, AC, TK helped with evoked response recordings. PP and TC helped with the morphological staining and analysis. All authors contributed to the writing and editing of the manuscript.

## Figure legends

**Suppl. Figure 1. Spontaneous L2/3 pyramidal neuron activity in Gabra1 KO mice. (A)** Field of view and ROI selection from RCaMP-expressing barrel cortex L2/3; (right panel) representative whole frame and individual traces from a whisker stimulation trial. **(B-B’’)** Average Ca^2+^ transient amplitude, duration and number of events in spontaneous trials in density plots and bar graphs (insets). **(C-C’)** Onset and decay times of Ca^2+^ transients in WT and *Gabra1* KO mice. The average was taken from all imaging sessions for individual neurons (N=459 neurons from 5 *Gabra1* KO mice, N=418 neurons from 4 WT mice). Statistics: linear mixed-effects models and Tukey post hoc tests. All bar graphs are represented as mean ± SEM. *p≤0.05, **p≤0.01, ***p≤0.001. **(D)** Immunohistochemistry of *Gabra1* WT and KO mice. Example images from L2/3 barrel cortex stained with VGAT, gephyrin and GABRA2 (α2) subunit. Scale bar: 25 μm. **(E-E’’’’)** Quantification of cluster density analyses of VGAT, gephyrin and GABRA2, and their colocalization. Statistics: t-test. Error bar: standard deviation. *p≤0.05

**Suppl. Figure 2. Spontaneous L2/3 pyramidal neuron activity in Gabra2 KO mice. (A)** Field of view and ROI selection from virus infected barrel cortex L2/3; (right panel) representative whole frame and individual traces from 1 whisker stimulation trial. **(B-B’’)** Average Ca^2+^ transient amplitude, duration and number of events in spontaneous trials in density plots and bar graphs (insets). **(C-C’)** Onset and decay times of Ca^2+^ transients in WT and *Gabra2* KO mice. The average was taken from all imaging sessions for individual neurons (N=370 neurons from 4 *Gabra2* KO mice, N=307 neurons from 3 WT mice). Statistics: linear mixed-effects models and Tukey post hoc tests. All bar graphs are represented as mean ± SEM. *p≤0.05, **p≤0.01, ***p≤0.001. **(D)** Immunohistochemistry of *Gabra2* WT and KO mice. Example images from L2/3 barrel cortex stained with VGAT, gephyrin and GABRA1 (α1) subunit. Scale bar: 25 μm. **(E-E’’’’)** Quantification of cluster density analyses of VGAT, gephyrin and GABRA1, and their colocalization. Statistics: t-test (unpaired), mean ± SD.

**Suppl. Figure 3. Changes in α1, α2, α3 GABA_A_R subunit expression in *Gabra1* and *Gabra2* KO mice.** Whole cell protein lysates were prepared from barrel cortices of WT & *Gabra1* KO littermates **(A)** and WT and *Gabra2* KO littermates **(B)** (5 mice per group), and Western blot (WB) analysis was performed for gephyrin, α3, and α1 or α2. Protein level was normalised to total actin level for further quantification. **(A’)** Quantification for gephyrin, α2, α3 protein expression changes between WT and *Gabra1* KO. **(B’)** Quantification for gephyrin, α1, α3 protein expression changes between WT and *Gabra2* KO. Statistics: t-test (unpaired). All bar graphs are represented as mean ± SD. *p≤0.05, **p≤0.01, ***p≤0.001.

**Suppl. Figure 4. Expression of RCaMP and eGFP_Cre and the neuronal activity in L2/3 pyramidal neuron *Gabra1*^*flox/flox*^ and *Gabra2*^*flox/flox*^ mice. (A, B)** Immunohistochemistry from *Gabra1*^*flox/flox*^ **(A)** and *Gabra2*^*flox/flox*^ **(B)** mice received viral injections of AAV6-CaMKIIa-RCaMP1.07 and AAV8-CaMKIIα-eGFP-Cre. The brain sections were stained for GABRA1 or GABRA2 to validate the effects of conditional knockout for *Gabra1* and *Gabra2*, respectively, in L2/3 pyramidal neurons. Scale bar: 25 μm; 10 μm for zoom-in images. **(C, D)** Example images of the field of view and ROI selection for Cre-negative and Cre-positive RCaMP-expressing pyramidal neurons of *Gabra1*^*flox/flox*^ **(C)** and *Gabra2*^*flox/flox*^ **(D)** mice. Right panel: example traces in a trial with whisker stimulation. Grey bar: whisker stimulation period. **(E, F)** Density distribution of amplitude, duration and number of events for spontaneous and stimulation-induced Ca^2+^ transients from RCaMP control, Cre-negative cells expressing RCaMP, Cre-positive cells expressing RCaMP in *Gabra1*^*flox/flox*^ **(E)** and in *Gabra2*^*flox/flox*^ **(F)** mice.

**Suppl. Figure 5. mIPSC and eIPSC recorded from gephyrin-DN-expressing L2/3 pyramidal neurons. (A)** Normal distribution was plotted and tested using QQ plot and D’Agostino-Pearson test. **(B)** logIEI plot for control and gephyrin variants. Statistics: One-way or Brown-Forsythe and Welch One way ANOVA followed by Tukey’s or Dunnet T3 multiple comparison tests. **(C)** Representative traces of evoked inhibitory postsynaptic currents (eIPSC) in a L2/3 pyramidal neuron of the barrel cortex of mice overexpressing control-eGFP and gephyrin-DN mutant. **(D)** Input/output curves and bar diagrams showing the eIPSC amplitude (pA) of L2/3 pyramidal neurons in GFP-expressing controls as compared to DN-expressing mutant mice for each stimulation paradigm (black, GFP: 4 mice, n=5; red, DN: 4 mice, n=4; Two-way ANOVA, F(8, 56) = 3.745, p = 0.0014; Post hoc Sidak’s multiple comparisons test, *p≤0.05). Data are reported as mean ± SEM.

**Suppl. Figure 6. Percentage of L2/3 pyramidal neurons infected with different AAV virus. (A)** Example images of barrel cortex L2/3 cells infected with AAV6-CaMKIIα-RCaMP1.07, AAV6-CaMKIIα-CreER^T2^ and AAV6-hSyn1-flex-gephyrin mutants. The neurons were stained for CaMKIIα and GFP. Scale bar: 20 μm. (B) Quantification of average expression of RCaMP-expressing neurons (72 %) and neurons co-expressing RCaMP and eGFP-gephyrin mutants (40 %) after normalization to total CaMKIIα-positive neurons.

**Suppl. Figure 7. Spontaneous activity changes in L2/3 pyramidal neurons expressing gephyrin variants. (A-A’’)** Average percentage changes after gephyrin-mutant expression for individual neurons are shown in bar graphs and the ratio are shown in violin plots to demonstrate the distributions for spontaneous Ca^2+^ transients. **(B)** Proportions of high- and lower-responding neurons before and after gephyrin-mutant expression *in vivo* selected based on Ca^2+^ transient amplitude. **(C)** Proportions of high- and lower-responding neurons before and after gephyrin-mutant expression selected based on number of events. eGFP, n=491 neurons; gephyrin-K148R, n=308 neurons; gephyrin-SSA, n=249 neurons; gephyrin-DN, n=204 neurons. The numbers on the pie charts are rounded to the nearest integer. Statistics: linear mixed-effects models and Tukey post hoc tests. All bar graphs are represented as mean ± SEM. *p≤0.05, **p≤0.01, ***p≤0.001.

