## Supplementary figures and images for "Differential impact of GABA_A_ receptors and gephyrin post-translational modifications on layer 2/3 pyramidal neuron responsiveness *in vivo*"

### Suppl. Figure 1

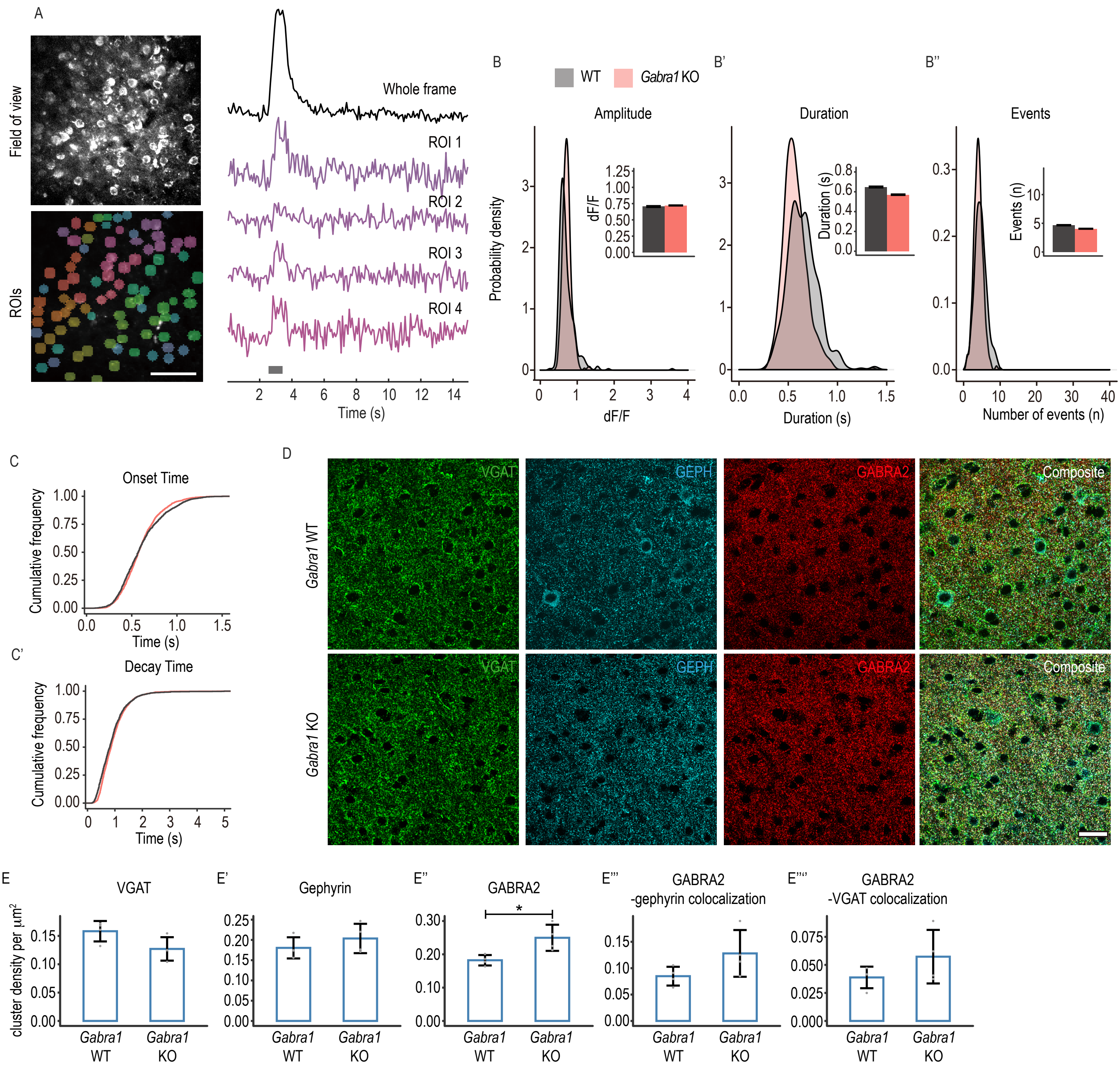


Supplementary figure 1.

### Suppl. Figure 2

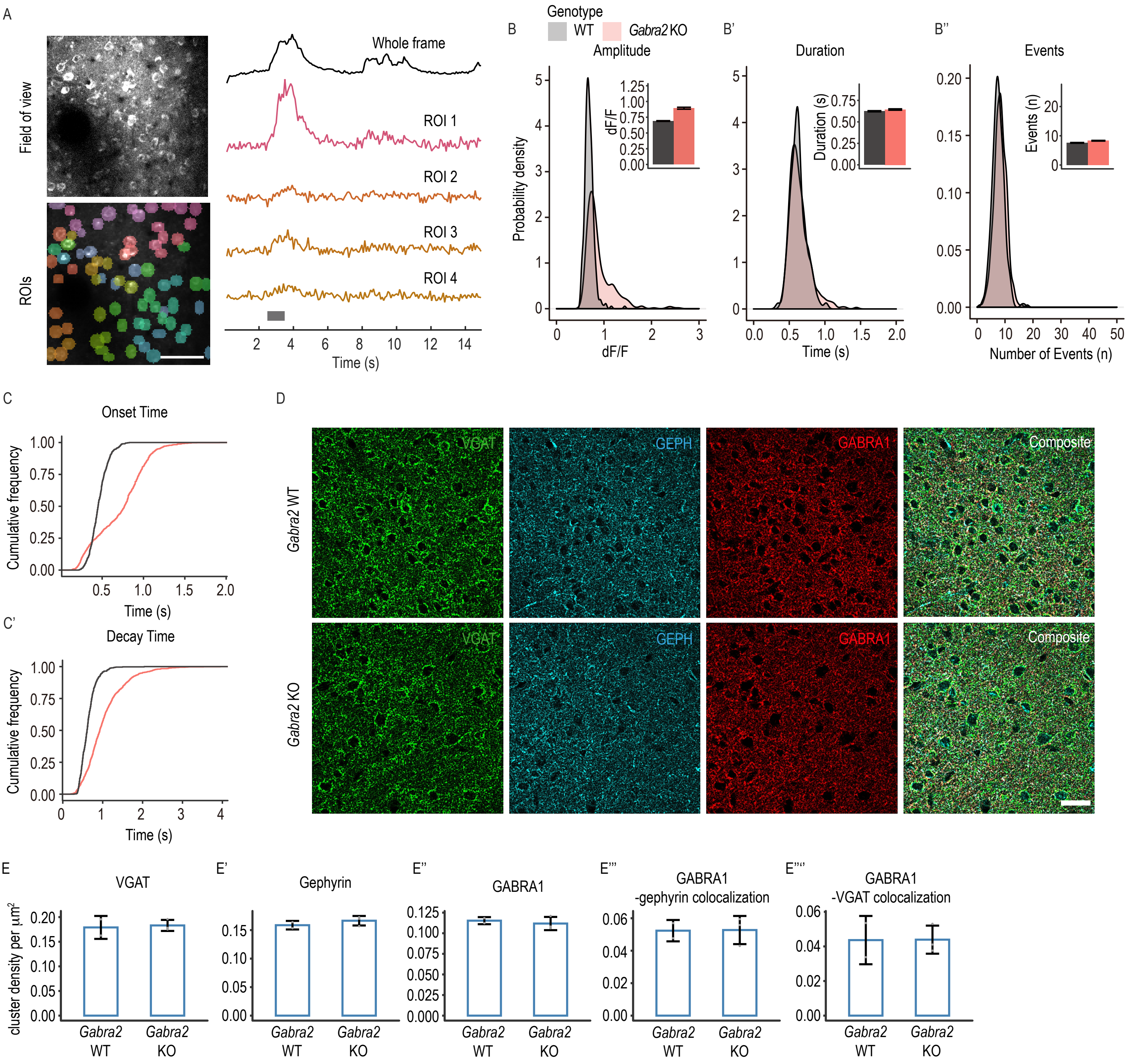


Supplementary figure 2.

### Suppl. Figure 3

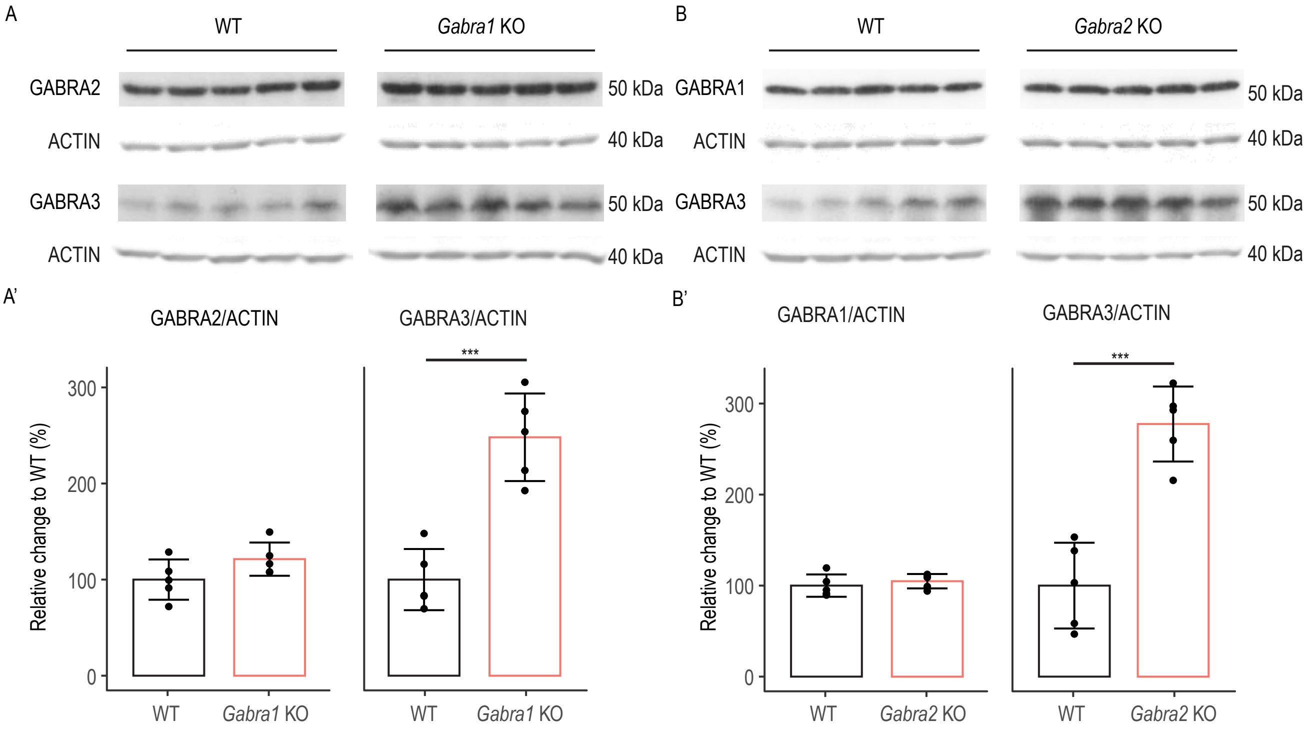


Supplementary figure 3.

### Suppl. Figure 4

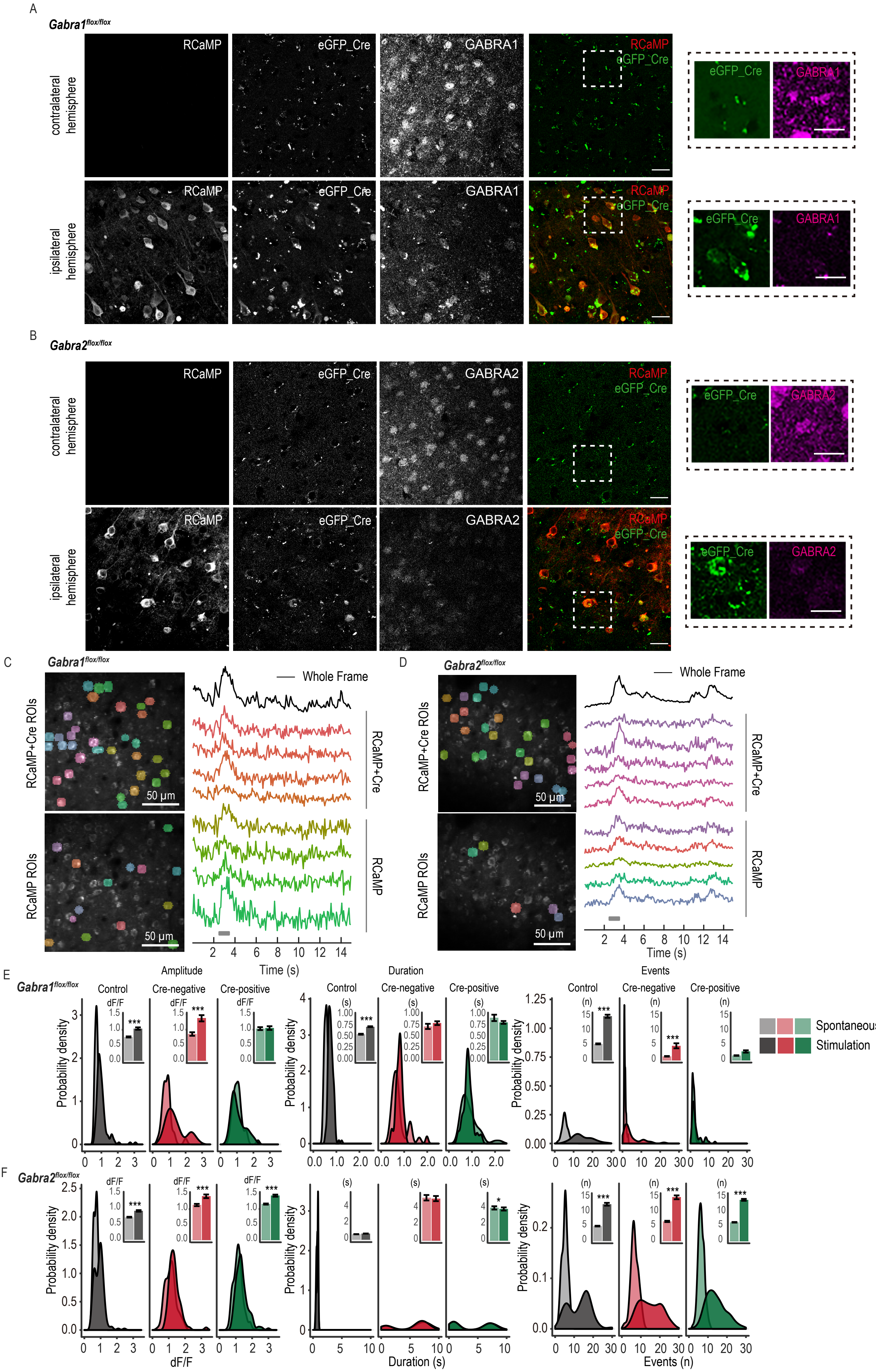


Supplementary figure 4.

### Suppl. Figure 5

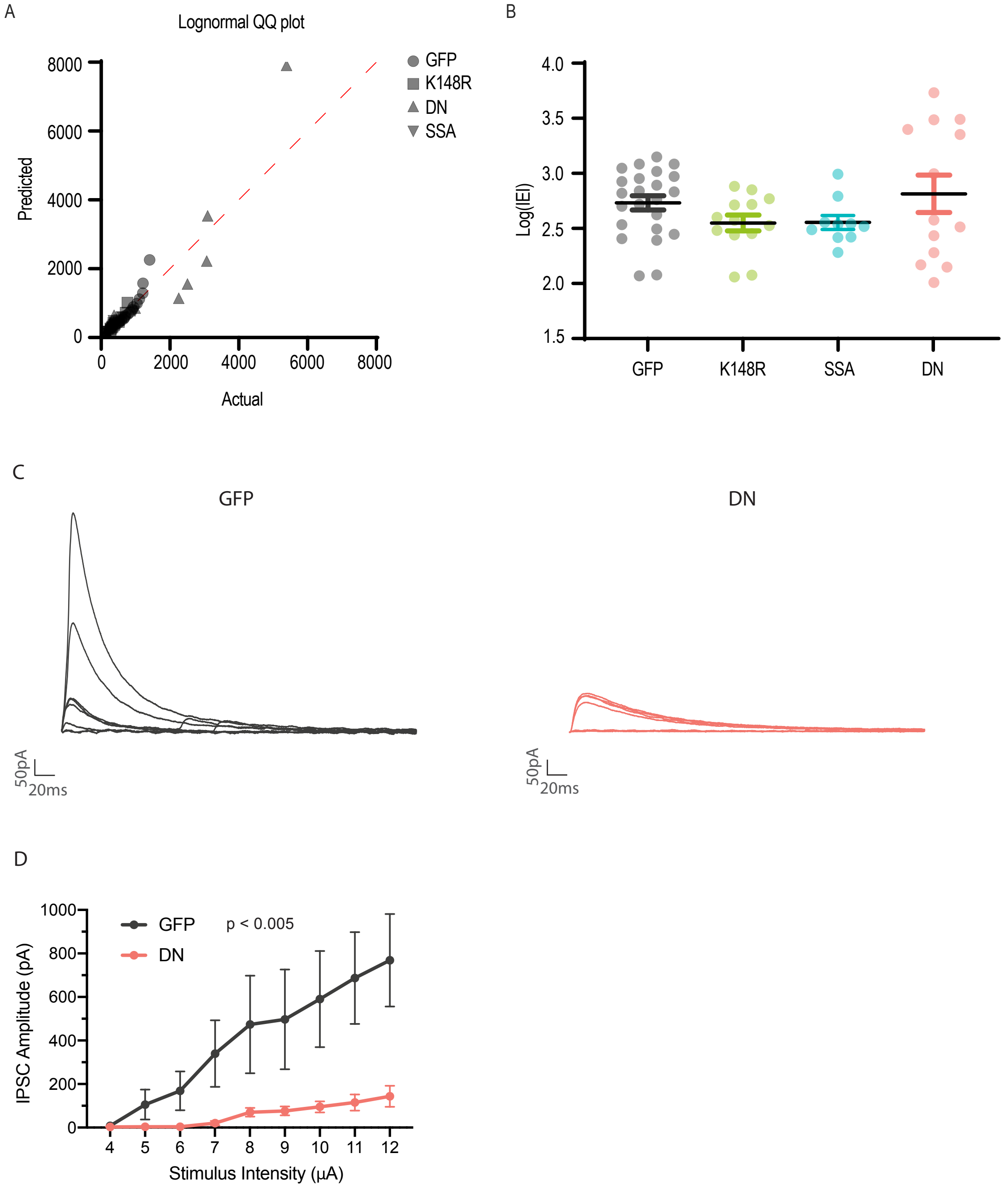


Supplementary Figure 5.

### Suppl. Figure 6

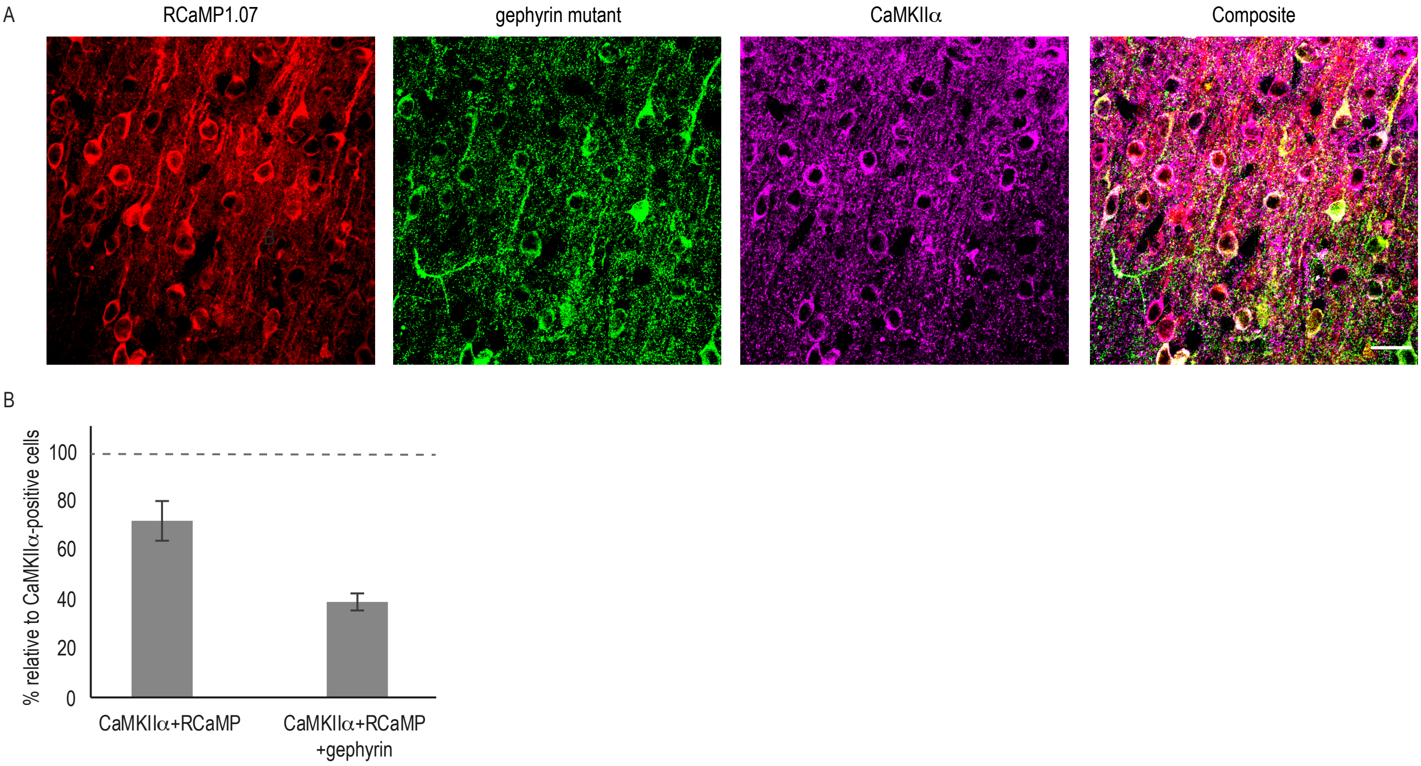


Supplementary figure 6.

### Suppl. Figure 7

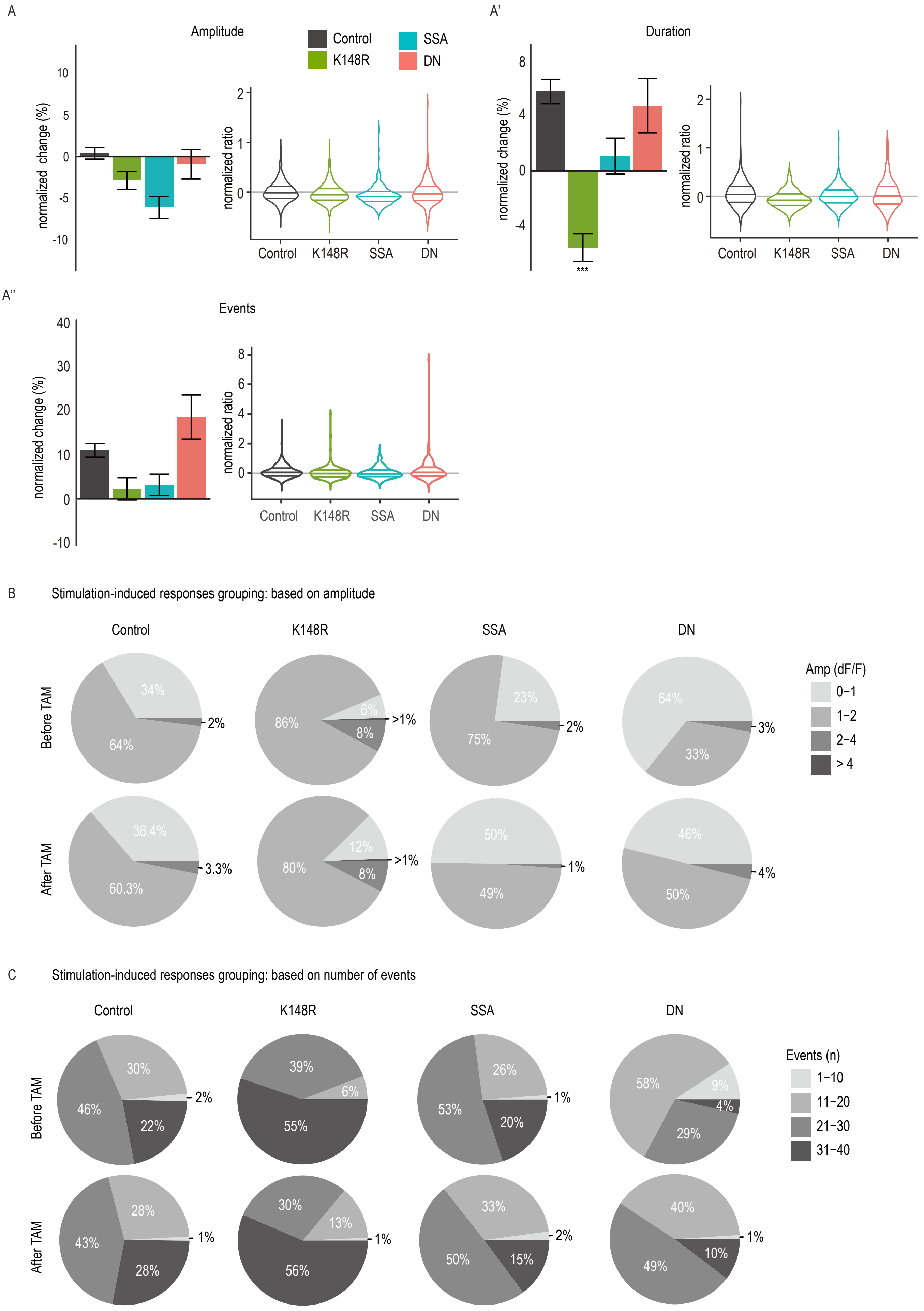


Supplementary figure 7.
